# A Meta-Analysis of fMRI Studies of Semantic Cognition in Children

**DOI:** 10.1101/2021.05.17.442947

**Authors:** Alexander Enge, Rasha Abdel Rahman, Michael A. Skeide

## Abstract

Our capacity to derive meaning from things that we see and words that we hear is unparalleled in other animal species and current AI systems. Despite a wealth of functional magnetic resonance imaging (fMRI) studies on where different semantic features are processed in the adult brain, the development of these systems in children is poorly understood. Here we conducted an extensive database search and identified 50 fMRI experiments investigating semantic world knowledge, semantic relatedness judgments, and the differentiation of visual semantic object categories in children (total *N* = 1,018, mean age = 10.1 years, range 4–15 years). Synthesizing the results of these experiments, we found consistent activation in the bilateral inferior frontal gyri (IFG), fusiform gyri (FG), and supplementary motor areas (SMA), as well as in the left middle and superior temporal gyri (MTG/STG). Within this system, we found little evidence for age-related changes across childhood and high overlap with the adult semantic system. In sum, the identification of these cortical areas provides the starting point for further research on the mechanisms by which the developing brain learns to make sense of its environment.

## 1. Introduction

The human capacity to retrieve meaning from words, phrases, and visual objects far exceeds the capacities of other animal species as well as all current state-of-the-art machine learning architectures. Functional magnetic resonance imaging (fMRI) has made it possible to map the brain areas underlying this capacity in the adult brain, showing that semantic information is processed in a distributed fashion across large parts of the cerebral cortex (Humphries et al., 2007; Huth et al., 2012, 2016; Liuzzi et al., 2020; Pulvermüller et al., 2009; Tyler et al., 2003). Most areas of this semantic system act largely amodal, that is, they show similar levels and patterns of activation regardless of whether the sensory input that is being processed comes from the visual domain or from the auditory domain (Deniz et al., 2019; Fairhall & Caramazza, 2013).

It is important, however, to interpret the findings from any individual fMRI study with caution: The generalizability of the patterns of brain activity that was observed may be limited to the respective task setting, stimuli, and population of participants under study (Yarkoni, 2021). Even when this is taken into account, the number of participants in a typical fMRI study is low (usually *N* ≤ 30). Therefore, many statistically significant peaks of activation may turn out to be spurious, capitalizing on chance fluctuations in the sample rather than genuinely task-related brain responses in the population (Button et al., 2013; Ioannidis, 2005; Thirion et al., 2007). One effective way of mitigating these two limitations is by statistically pooling the results from individual fMRI experiments on a given topic into a meta-analysis. This can be done in an image-based fashion, using the statistical parametric maps from the original experiments, or in a coordinate-based fashion, using only the peak coordinates. Despite evidence that inferences from the image-based approach are more precise (Salimi-Khorshidi et al., 2009), this approach remains difficult to implement since statistical maps for most fMRI experiments are still not being shared (Poline et al., 2012). In contrast, peak coordinates are routinely reported in research articles, oftentimes making the coordinate-based approach the only feasible one in practice (Samartsidis et al., 2017).

While many of such coordinate-based meta-analyses have been conducted for fMRI studies investigating semantic cognition in adults (Binder et al., 2009; Cocquyt et al., 2019; Ferstl et al., 2008; Jackson, 2021; Noonan et al., 2013; Rodd et al., 2015; Vigneau et al., 2006; Visser et al., 2010; Wu et al., 2012), none is available as of yet to complement this effort from a developmental perspective. One reason for this may be that children are more difficult to recruit and scan than adult participants. They often require specialized equipment, additional training sessions (e.g., familiarizing them with a mock MRI scanner), and frequently the disposal of volumes or entire runs due to excessive motion or inattentiveness. Nevertheless, a number of studies have successfully used fMRI to investigate the development of semantic processing, providing preliminary evidence for how and when the neurobiological architecture for processing meaning comes about during childhood. In these studies, children were typically scanned while probing their semantic world knowledge (e.g., by asking the child to name an object after hearing its description or to decide if a certain word refers to something animate or inanimate; e.g., Balsamo et al., 2006), their judgements of the semantic relatedness between concepts (e.g., by asking the child if two sequentially presented words or pictures were related to one another or not; e.g., Chou et al., 2019), or their viewing of different semantic categories of visual objects (e.g., by having the child perform a visual detection task while passively viewing images of human faces, tools, and scenes; e.g., Scherf et al., 2007). At this point, synthesizing these heterogeneous efforts meta-analytically is becoming important (a) to distinguish between consistent and potentially spurious findings, (b) to identify the similarities and differences between different aspects of semantic cognition (i.e., between different task categories), and (c) to identify differences in the semantic system between children and adults.

Here we conducted such a coordinate-based meta-analysis of the currently available fMRI experiments probing semantic cognition in children. Based on a systematic search of the literature using online databases, we sought to identify a wide range of fMRI studies, covering different aspects of semantic cognition and a broad age range from early childhood until the beginning of adolescence. We hypothesized that general semantic cognition would be associated with consistent activation in many of the same areas that have been found during semantic processing in adults (see, e.g., Binder et al., 2009; Jackson, 2021), namely the left inferior frontal gyrus (IFG; especially the pars triangularis and pars orbitalis), the left middle temporal gyrus (MTG) and anterior temporal lobe (ATL), as well as regions known to be sensitive to the differences between semantic visual object categories, such as the fusiform gyrus (FG) and lateral occipital complex (LOC). In children, we expected to identify additional clusters of consistent activation in the right-hemispheric homologues of these regions. This is because the left-lateralization of the language comprehension network, despite being present from newborn age onwards, continues to fully develop until early adolescence (especially in the IFG; Berl et al., 2014; Enge et al., 2020; Holland et al., 2007).

## 2. Materials and Methods

### 2.1 Literature Search

The search terms “(child OR children OR childhood OR pediatric) AND (brain mapping OR brain scan OR functional magnetic resonance imaging OR functional MRI OR fMRI OR neuroimaging) AND (semantics OR category OR categorization OR conceptual knowledge OR semantic knowledge OR semantic memory OR semantic feature OR semantic category OR semantic categorization OR semantic comprehension OR visual semantics OR visual categorization OR object categorization)” were entered into three online databases (PubMed/MEDLINE, PsycInfo, and Scopus). As of July 2020, this search yielded a total of 1,095 articles. Of these, 895 remained after removing duplicate articles and were subsequently evaluated for eligibility (see Figure 1). We pre-specified ten inclusion criteria, ensuring that all articles to be included (1) were written in English, (2) reported original results from a group study (excluding review articles, meta-analyses, surveys, and case studies), (3) tested at least one group of children with a mean age of 3–12 years (range 3–15 years), (4) tested a typically developing, non-clinical sample (including healthy control groups from clinical studies), (5) performed task-based fMRI (excluding resting-state fMRI and other imaging modalities), (6) had children engage in a task probing semantic cognition (i.e., semantic world knowledge, semantic relatedness, or visual object semantics), (7) analyzed the fMRI data within the framework of the general linear model (GLM), (8) applied the same statistical threshold across the whole brain (excluding ROI analyses and partial brain coverage; Müller et al., 2018), (9) reported results as peak coordinates in standard space (Talairach or MNI), and (10) reported peaks for the within-group contrast of two semantic conditions and/or one semantic and one control condition.

**Figure 1.**
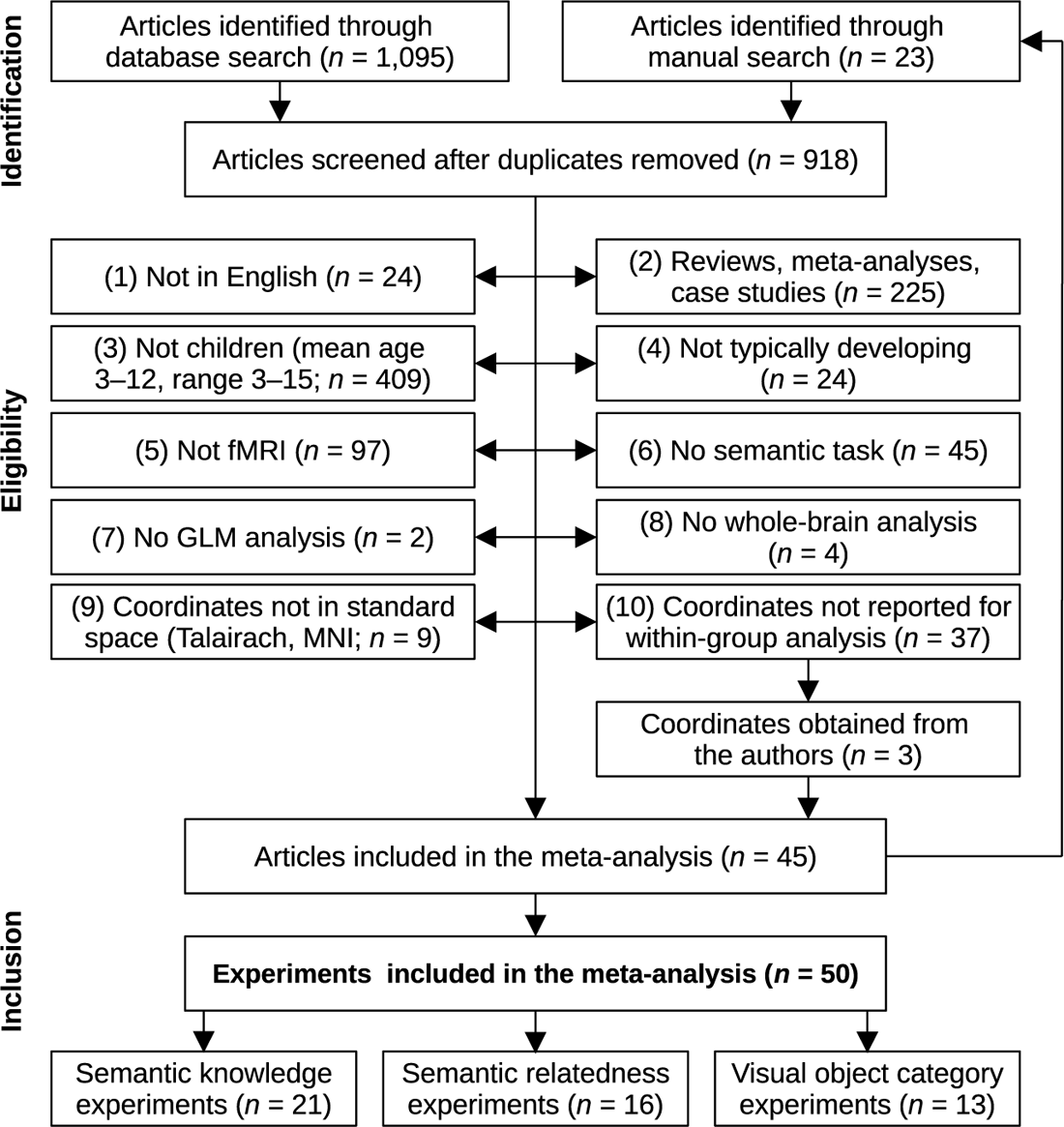
Literature Search and Selection Workflow

Initially, this led to the inclusion of 34 articles. We consulted the introduction and reference sections of these articles as well as relevant review papers on children’s semantic and language processing (Antonucci & Alt, 2011; Barquero et al., 2014; Enge et al., 2020; Leach & Holland, 2010; Martin et al., 2015; O’Shaughnessy et al., 2008; Sachs & Gaillard, 2003; Schlaggar & McCandliss, 2007; Skeide & Friederici, 2016; Weiss-Croft & Baldeweg, 2015) to identify additional articles not covered by our database search. Following this procedure, we identified 23 additional articles, seven of which fulfilled all inclusion criteria.

In addition to those articles fulfilling all ten inclusion criteria, 37 articles met all but the last criteria—that is, the relevant within-group peak coordinates were not reported in the published article. In these cases, we contacted the corresponding authors to request the missing information. This led to the inclusion of three additional articles, resulting in a total of 45 articles being included in the meta-analysis.

Whenever one of these articles reported multiple contrasts based on the same sample of children, the coordinates from all of these contrasts were treated as a single experiment (note that *experiment* is the term we use whenever we refer to our primary unit of analysis; Turkeltaub et al., 2012). This is considered good practice in order to minimize within-group effects and avoid inflating the number of independent data points included in the meta-analysis (Eickhoff et al., 2012; Turkeltaub et al., 2012). Conversely, whenever an article reported multiple contrasts from two or more independent samples of children, these were treated as separate experiments. This led to a final meta-analytic sample of *n* = 50 experiments. While each of these experiments targeted some aspect of general semantic cognition, they could further be subdivided into more homogeneous groups of experimental tasks, probing (a) semantic world knowledge (e.g., naming an object after hearing its description; *n* = 21), (b) semantic relatedness (e.g., hearing two words and deciding if they are related or not; *n* = 16), and (c) visual semantic object categories (e.g., passively viewing faces as compared to other visual stimuli; *n* = 13). The experiments belonging to these three task categories as well as the entire data set (probing general semantic cognition across all task categories) were meta-analyzed using two different algorithms: activation likelihood estimation and seed-based *d* mapping.

### 2.2 Activation Likelihood Estimation

Activation likelihood estimation (ALE) is the most frequently used algorithm to perform coordinate-based meta-analyses of neuroimaging experiments (Acar et al., 2018). It estimates the degree to which peak coordinates taken from independent MRI experiments, all investigating the same task and/or participant population, spatially converge to form non-random clusters of activation (Eickhoff et al., 2009, 2012; Turkeltaub et al., 2002). To this end, the algorithm first recreates a modeled activation map for each of the input experiments. All voxels for which the experiment reports a peak coordinate are assigned a value of 1, whereas all of the other voxels within the gray matter mask are assigned a value of 0. Because these peaks entail spatial uncertainty and are assumed to be part of larger clusters of activation, their values are smoothed across the neighboring voxels by convolving them with a Gaussian kernel. The width of this kernel is chosen to be inversely proportional to the sample size of the experiment, reflecting the fact that larger sample sizes provide stronger evidence for the true location of any peak of activation (Eickhoff et al., 2009). When two (or more) peaks in the experiment are reported in close proximity of one another, the voxels at which their Gaussians overlap are assigned the maximum—rather than the sum—of their respective values.

This prevents any meta-analytic cluster from receiving artificially high likelihood values merely because a large number of subpeaks has been reported for this cluster in the original article.

The modeled activation maps of the individual experiments are then combined into a single meta-analytic map by assigning an ALE value to each voxel. This ALE value is computed as the union of the modeled activation values for this voxel across the modeled activation maps for all experiments (Acar et al., 2018; Eickhoff et al., 2012):

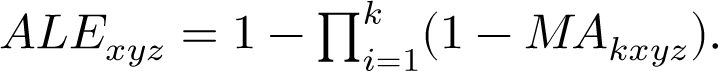

This hierarchical procedure treats the included experiments as a random subsample of all possible experiments and therefore allows the generalization across the population of possible fMRI experiments on the topic of interest (Eickhoff et al., 2009). The statistical significance of these voxel-wise ALE values is determined by comparing them to an analytically derived null distribution as described by Eickhoff et al. (2012). To correct for multiple comparisons, a cluster-level family-wise error (FWE) correction procedure has been shown to offer an excellent trade-off between control over the Type I error rate and statistical power (Eickhoff et al., 2016).

For the present analysis, these steps were performed using the NiMARE package (Version 0.0.7; Salo et al., 2021) in Python (Version 3.8.8; Van Rossum & Drake, 2009). If necessary, coordinates were transformed from Talairach to MNI space using the icbm2tal transform function (Lancaster et al., 2007). The modeled activation maps were rendered in MNI152 space at 2×2×2 mm resolution (Fonov et al., 2011). For statistical thresholding, a voxel-level cluster-forming threshold of *p* < .001 (uncorrected) and a cluster-level threshold of *p* < .01 (FWE-corrected) was used. This cluster-level threshold was determined by comparing the observed cluster size to an empirical distribution built from 1,000 iterations of drawing random peak locations from the gray matter template and recording the maximal cluster size. The Nilearn package (Version 0.7.1; Abraham et al., 2014) was used for image processing and plotting while the anatomic automatic labeling atlas (AAL2; Rolls et al., 2015) as implemented in the AtlasReader package (Version 0.1.2; Notter et al., 2019) was used for anatomical labelling.

### 2.3 Seed-Based *d* Mapping

While ALE estimates the spatial convergence of reported activation peaks, an alternative approach is to use the effect sizes (if available) of these peaks to infer a meta-analytic effect size for each gray matter voxel. This approach more closely resembles traditional meta-analyses of behavioral or clinical outcomes and is used by the seed-based *d* mapping (SDM) algorithm (Albajes-Eizagirre, Solanes, Vieta, et al., 2019). In short, it determines lower and upper bounds for possible effect size images based on the peak coordinates and their reported effect sizes (*t* scores or *z* scores). Then, a meta-analytic method correcting for non-statistically significant unreported effects (MetaNSUE; Albajes-Eizagirre, Solanes, & Radua, 2019) is used to infer the most plausible effect size and its standard error based on multiple imputations of censored information. Subsequently, all imputed data sets are meta-analyzed separately and combined using Rubin’s rules. For statistical thresholding, the resulting meta-analytic map is FWE-corrected by comparing the voxel-wise observed effect size against an empirical null distribution of effect sizes built from random permutations.

The SDM algorithm was used on the same experiments and peak coordinates as for the ALE analysis but adding, if available, their reported *t* scores or *z* scores (the latter being converted to *t* scores with *df* = *n_children_* - 1). The data were preprocessed with the SDM-PSI software (Version 6.21, https://www.sdmproject.com), using its default gray matter correlation template with a voxel size of 2×2×2 mm and a Gaussian smoothing kernel (anisotropy α = 1.0, FWHM = 20 mm). This effect-size based SDM algorithm made it possible (a) to probe the robustness of the ALE results against a change in the meta-analytic approach and (b) to statistically control for systematic differences between the included experiments by means of a covariate analysis. Note that the latter type of analysis is impossible in an approach like ALE because it disregards the effect sizes of the reported peaks and instead treats them as binary. Two separate models were computed to meet these two objectives: (a) a mean-based meta-analysis without any covariates or predictors (as in the ALE analysis) and (b) a mean-based meta-analysis controlling for four different experiment-level confounds (mean age of the children [mean-centered continuous predictor], presentation modality [0 = visual, 1 = audiovisual, 2 = auditory and visual in separate blocks, 3 = auditory], response modality [0 = no response, 1 = manual response, 2 = covert speech, 3 = overt speech], and data analysis software [0 = SPM, 1 = FSL, 2 = other]. Both models were estimated with 50 random imputations and statistical thresholding was performed using a voxel-level FWE-corrected threshold of *p* < .001 and a cluster extent threshold of *k* > 25 connected voxels (200 mm^3^).

### 2.4 Differences Between Semantic Task Categories

Beyond mapping the cortical network associated with general and task-specific semantic cognition in children, a meta-analytic subtraction analysis (Laird et al., 2005) was carried out to test for reliable differences between the three different task categories (i.e., knowledge, relatedness, and objects). For this type of analysis, one ALE map (e.g., the map for the semantic knowledge experiments) was subtracted from another ALE map (e.g., the combined map for the semantic relatedness and visual object category experiments). The resulting map of difference scores was then compared against an empirical null distribution of such difference maps, obtained from randomly reshuffling the original experiments 20,000 times into new groups. Voxels with *p* < .001 (uncorrected) as compared to this null distribution and forming clusters of at least 25 connected voxels (200 mm^3^) were considered as showing reliable differences between task categories. In addition, a meta-analytic conjunction analysis was performed to identify areas where cognitive processing was shared across all three task categories. This was done by taking the minimum ALE value at each voxel across all three task-specific ALE maps—but only for those voxels that were statistically significant in each of the three (Nichols et al., 2005).

### 2.5 Age-Related Changes

The same approach as just described was used to compare semantic cognition in older versus younger children. To this end, the original sample was split into equally sized groups at the median of the (mean) sample ages across experiments. These two groups of experiments were compared using the same subtraction procedure and statistical threshold as for the semantic task categories.

Additionally, we also tested for a linear influence of age by means of a meta-regression using SDM (see Section 2.3). In this linear model, the outcome of interest was not the voxel-wise effect size across experiments (as in the main SDM analysis) but those voxels whose effect size showed significant covariation with the (mean) age of the sample(s) of children contributing to it.

### 2.6 Comparison With Semantic Cognition in Adults

The meta-analytic results of semantic cognition in children were also compared to semantic cognition in adults as reported in a recent meta-analysis on semantic control (Jackson, 2021). To this end, we recreated their ALE analysis of general semantic cognition in adults (*n* = 415 experiments; see their Figure 3 and Table 3) and compared it to our child-specific ALE analysis by means of a meta-analytic subtraction and conjunction analysis as described above (see Section 2.4).

### 2.7 Evaluation of Robustness

Meta-analyses reflect the state of the published literature on a given topic and are therefore subject to the same biases as the original studies (e.g., small sample bias, selective reporting, file drawer problem). For behavioral and clinical meta-analyses, a range of standard tools has been developed to assess the risk of these biases as well as the robustness of the meta-analytic results against them. Some but not all of these tools can be carried over to the meta-analysis of neuroimaging data (Acar et al., 2018). For instance, it is possible to assess the degree to which the meta-analytic results depend on any individual study, which may or may not have reported false positive findings (e.g., due to low statistical power; Button et al., 2013; Ioannidis, 2005; Thirion et al., 2007). This can be done by recomputing the original meta-analysis as many times as there are experiments included, each time leaving out one of these experiments. This leave-one-out analysis (also called jackknife analysis) reveals if any of the observed meta-analytic clusters critically depends on a single influential experiment or if it is robust against a false positive experiment in the sample.

Publication bias may not only manifest itself in the form of published experiments reporting false positive effects but also in the form of experiments not getting published when failing to obtain statistically significant effects. Because of this “file drawer” problem, there are up to approximately 30 unpublished neuroimaging experiments with null effects (i.e., reporting zero significant peaks) per 100 published experiments (Samartsidis et al., 2020). While these cannot be factored into the meta-analysis directly, the simulation of imaginary file drawers with different numbers of null experiments is informative regarding the robustness of the results against this type of bias. In this context, the fail-safe N (FSN) metric has been defined as the number of null experiments that can be added to the original meta-analysis without rendering its meta-analytic effect size statistically non-significant (Rosenthal, 1979). If FSN exceeds the upper bound of experiments that can realistically be expected to be inside the file drawer, one can conclude that the file drawer problem does not suffice to explain the meta-analytic result. This logic can be extended to meta-analyses of neuroimaging studies by simulating null experiments with peaks of activation at random rather than spatially converging locations across the brain (Acar et al., 2018). Such null experiments were simulated as to resemble the original experiments in terms of their individual sample sizes and numbers of reported peak coordinates but had their peak locations drawn randomly from all possible voxels within the gray matter template. They were then added iteratively to the original experiments. At each step, the ALE analysis was repeated, recording for every voxel if it had remained part of a statistically significant cluster or not. This was repeated up to a maximum of five times the number of experiments in the original sample (e.g., FSN_max_ = 150 for our main analysis with *n* = 50 experiments). The whole procedure was repeated for 10 different (random) file drawers of null experiments.

Both of these approaches (leave-one-out and fail-safe N analysis) were performed separately for our main ALE analysis (including all 50 experiments) as well as for the task category-specific sub-analyses. They were expected to be especially informative for the category-specific analyses because the low number of experiments in these sub-analyses might have reduced the robustness of the meta-analytic results (Eickhoff et al., 2016).

## 3. Results

### 3.1 Literature Search

As of July 2020, a total of 45 articles reporting 50 independent fMRI experiments of semantic cognition in children were obtained by searching online literature databases (see Figure 1 and Table 1). These experiments could be grouped further into experiments probing semantic world knowledge (*n* = 21), semantic relatedness judgments (*n* = 16), and the discrimination of visual semantic object categories (*n* = 13). Together, they comprise fMRI data of 1,018 children (*m* = 20.4 per experiment, *md* = 15.5, range 5–67; see Figure 2) with a mean age of 10.1 years (range of mean ages 5.5–12.8 years, total age range 4–15 years). According to the original articles, 54.4% of these children were boys (45.6% were girls) and 98.4% were right-handed (1.6% were left-handed). From these experiments, a total of 687 peaks of activation were reported (*m* = 13.7 per experiment, *md* = 12, range 1–47) and entered into the meta-analysis. Of these peaks, 400 (58.2%) were in the left hemisphere (*x*_MNI_ < 0), indicating a slight degree of lateralization. There were no reliable associations across experiments between sample size, the mean age of the children under study, and the number of peaks reported (see Figure 2). Additional descriptive information about the experiments can be found in the appendix.

**Figure 2.**
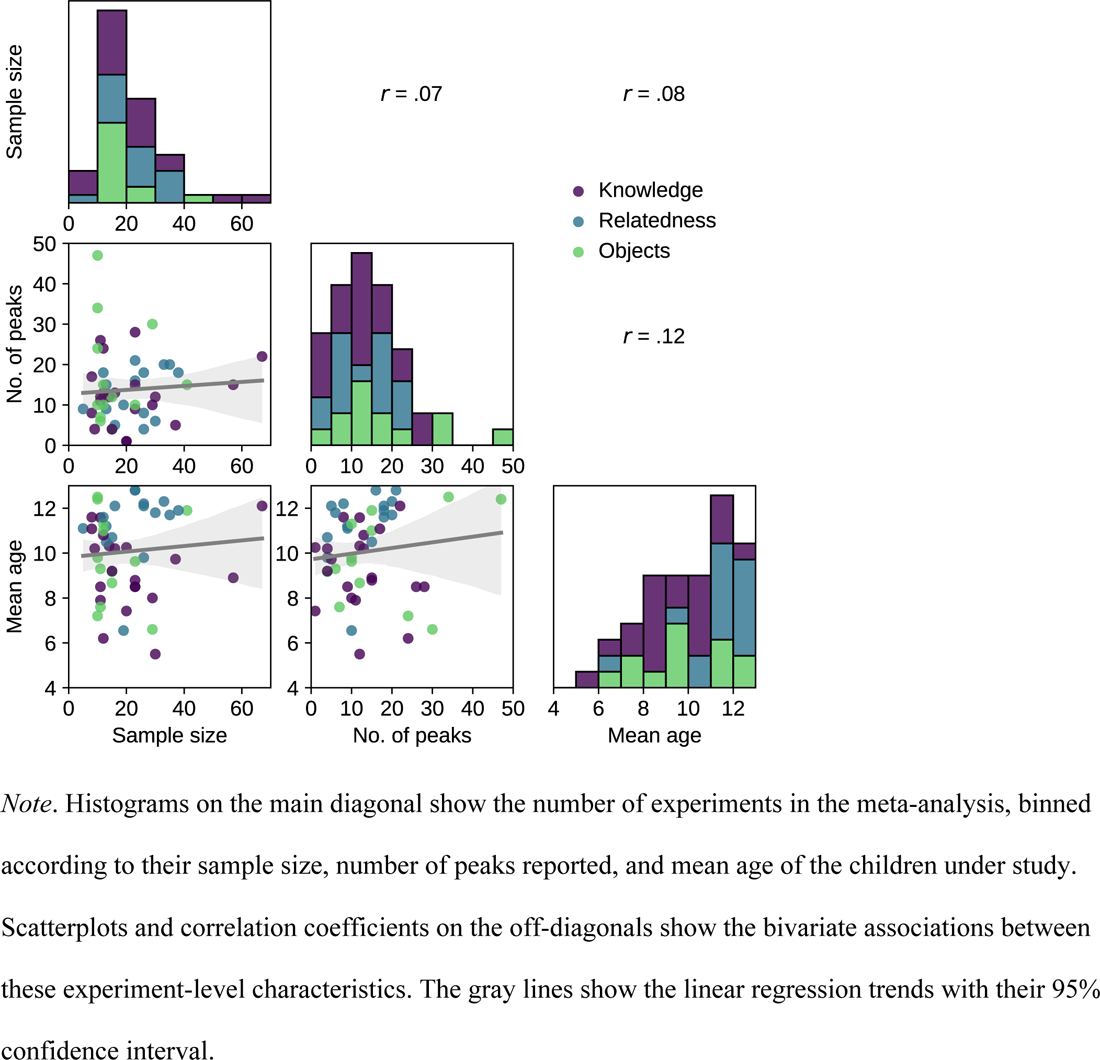
Distributions and Bivariate Associations of Experiment-Level Characteristics

**Table 1.** Experiments Included in the Meta-Analysis

| # | Author(s) (year) | *n* | Sex<br>(F/M) | Hand.<br>(R/L) | Mean age<br>(SD) | Age<br>range | Semantic task | Task<br>category | No. of<br>peaks |
| --- | --- | --- | --- | --- | --- | --- | --- | --- | --- |
| 1 | Arnoldussen (2006) | 11 | NA | NA | 11.6 (1.0) | 9–13 | Picture–word matching vs. shape matching | Knowledge | 12 |
| 2 | Arnoldussen (2006) | 11 | NA | NA | 7.9 (0.6) | 7–8 | Picture–word matching vs. shape matching | Knowledge | 11 |
| 3 | Aylward et al. (2005) | 11 | 4/7 | 10/1 | 9.3 (0.9) | 8–10 | Viewing faces vs. viewing houses | Objects | 6 |
| 4 | Backes et al. (2002) | 8 | 0/8 | 8/0 | 11.6 (0.7) | 11–12 | Animal/no-animal judgment vs. fixation | Knowledge | 8 |
| 5 | Balsamo et al. (2002) | 11 | 7/4 | 11/0 | 8.5 (0.9) | 7–9 | Naming after description vs. rest | Knowledge | 26 |
| 6 | Balsamo et al. (2006) | 23 | 13/10 | 22/1 | 8.5 (1.5) | 5–10 | Category–word judgment vs. tone detection | Knowledge | 9 |
| 7 | Bauer et al. (2017) | 14 | 7/7 | 14/0 | 10.3 (0.9) | 8–11 | Object size or animacy judgment vs. fixation | Knowledge | 12 |
| 8 | Berl et al. (2014) | 57 | 26/31 | 57/0 | 8.9 (NA) | 4–12 | Definition–word matching vs. tone detection | Knowledge | 15 |
| 9 | Booth et al. (2001) | 5 | 0/5 | 5/0 | 11.1 (NA) | 9–12 | Word relatedness judgement vs. tone/symbol matching | Relatedness | 9 |
| 10 | Booth et al. (2003) | 15 | 8/7 | 15/0 | 10.7 (NA) | 9–11 | Word relatedness judgement vs. tone/symbol matching | Relatedness | 4 |
| 11 | Booth et al. (2007) | 13 | 4/9 | 13/0 | 10.5 (2.2) | 9–15 | Word relatedness judgement vs. false font matching | Relatedness | 15 |
| 12 | Brauer & Friederici (2007) | 12 | 8/4 | 12/0 | 6.2 (NA) | 5–6 | Semantic sentence acceptability judgment vs. rest | Knowledge | 24 |
| 13 | Cao et al. (2008) | 13 | 6/7 | 13/0 | 11.2 (NA) | 9–12 | Word relatedness judgement vs. symbol string matching | Relatedness | 9 |
| 14 | Chou et al. (2006) | 35 | 22/13 | 35/0 | 11.7 (2.1) | 9–15 | Word relatedness judgment vs. color change detection | Relatedness | 20 |
| 15 | Chou et al. (2006) | 26 | NA | 26/0 | 12.1 (2.0) | 9–15 | Word relatedness judgment vs. color change detection | Relatedness | 18 |
| 16 | Chou et al. (2009) | 33 | 16/17 | 33/0 | 12.3 (1.8) | 8–15 | Word relatedness judgment vs. non-character matching | Relatedness | 20 |
| 17 | Chou et al. (2019) | 16 | 5/11 | 16/0 | 12.1 (1.4) | 10–14 | Word relatedness judgement (multiple contrasts) | Relatedness | 5 |
| 18 | Corbett et al. (2009) | 15 | 2/13 | NA | 9.2 (1.4) | 8–12 | Sequential matching of faces vs. things | Objects | 4 |
| 19 | Dekker et al. (2014) | 10 | 2/8 | 10/0 | 9.8 (0.4) | 9–10 | Categorical one-back task (multiple categories) | Objects | 10 |
| 20 | Dekker et al. (2014) | 11 | 4/7 | 11/0 | 7.6 (0.4) | 7–8 | Categorical one-back task (multiple categories) | Objects | 7 |
| 21 | Fan et al. (2020) | 26 | 14/12 | 26/0 | 9.8 (1.5) | 8–12 | Word relatedness judgment vs. visual cue detection | Relatedness | 4 |
| 22 | Gaillard et al. (2001) | 9 | 4/5 | 9/0 | 10.2 (NA) | 7–13 | Naming after description vs. dot pattern viewing | Knowledge | 4 |
| 23 | Gaillard et al. (2003) | 16 | 9/7 | 16/0 | 10.2 (NA) | 7–14 | Word generation to category names vs. rest | Knowledge | 13 |
| 24 | Horowitz-Kraus et al. (2015) | 23 | 15/8 | 23/0 | 8.5 (0.8) | 7–9 | Sentence–picture matching vs. word–picture matching | Knowledge | 28 |
| 25 | Kersey et al. (2016) | 29 | 12/17 | NA | 6.6 (NA) | 4–8 | Picture matching with tools vs. other categories | Objects | 30 |
| 26 | Krishnan et al. (2015) | 37 | 19/18 | 37/0 | 9.7 (NA) | 7–12 | Picture matching vs. saying “silly” | Knowledge | 5 |
| 27 | Krishnan et al. (2021) | 67 | NA | NA | 12.1 (1.7) | 10–15 | Verb generation to pictures vs. rest | Knowledge | 22 |
| 28 | Lee et al. (2011) | 23 | 11/12 | 23/0 | 12.8 (1.5) | 10–15 | Word relatedness judgment vs. word–tone matching | Relatedness | 21 |
| 29 | Lee et al. (2011) | 23 | 11/12 | 23/0 | 12.8 (1.5) | 10–15 | Word relatedness judgment vs. non-character matching | Relatedness | 16 |
| 30 | Lee et al. (2016) | 30 | 14/16 | 30/0 | 11.8 (1.9) | 8–15 | Word relatedness judgment vs. non-character matching | Relatedness | 6 |
| 31 | Libertus et al. (2009) | 15 | 7/8 | NA | 8.7 (NA) | 8–9 | Two-back task with faces vs. other categories | Objects | 12 |
| 32 | Liebig et al. (2017) | 41 | 18/23 | 41/0 | 11.9 (NA) | 9–13 | Animacy judgment vs. visual symbol judgment | Objects | 15 |
| 33 | Mathur et al. (2020) | 19 | 11/8 | 19/0 | 6.6 (NA) | 5–7 | Word–picture relatedness judgment vs. symbol string matching | Relatedness | 10 |
| 34 | Meyler et al. (2008) | 12 | 9/3 | 12/0 | 10.8 (0.4) | NA | Semantic sentence acceptability judgment vs. fixation | Knowledge | 13 |
| 35 | Monzalvo et al. (2012) | 23 | 11/12 | NA | 9.6 (0.5) | 8–10 | Target detection while viewing faces/houses vs. other categories | Objects | 10 |
| 36 | Moore-Parks et al. (2010) | 23 | 12/11 | 23/0 | 8.8 (1.1) | 7–10 | Definition–word matching vs. reversed speech | Knowledge | 15 |
| 37 | Okamoto et al. (2017) | 12 | 1/11 | 11/1 | 11.3 (1.3) | NA | Viewing bodies/faces vs. other categories | Objects | 10 |
| 38 | Passarotti et al. (2003) | 12 | 8/4 | 12/0 | NA (NA) | 10–12 | Face matching vs. detecting scrambled faces | Objects | 15 |
| 39 | Schafer et al. (2009) | 26 | 15/11 | 24/2 | 12.2 (0.4) | NA | Word relatedness judgment vs. visual symbol judgment | Relatedness | 8 |
| 40 | Scherf et al. (2007) | 10 | 4/6 | 10/0 | 12.5 (1.0) | 11–14 | Viewing movies of faces/places/objects vs. other categories | Objects | 34 |
| 41 | Scherf et al. (2007) | 10 | 4/6 | 10/0 | 7.2 (1.0) | 5–8 | Viewing movies of faces/places/objects vs. other categories | Objects | 24 |
| 42 | Scherf et al. (2010) | 10 | 0/10 | 10/0 | 12.4 (1.3) | 10–14 | Viewing movies of faces/places/objects vs. other categories | Objects | 47 |
| 43 | Siok et al. (2004) | 8 | 4/4 | 8/0 | 11.1 (NA) | 10–12 | Chinese character judgment vs. fixation | Knowledge | 17 |
| 44 | Skeide et al. (2014) | 20 | 8/12 | 20/0 | 10.3 (NA) | 9–10 | Picture–sentence matching (plausible vs. implausible) | Knowledge | 1 |
| 45 | Skeide et al. (2014) | 20 | 11/9 | 20/0 | 7.4 (NA) | 6–7 | Picture–sentence matching (plausible vs. implausible) | Knowledge | 1 |
| 46 | Szaflarski et al. (2006) | 29 | 15/14 | NA | NA (NA) | 5–11 | Verb generation to nouns vs. finger tapping to tones | Knowledge | 10 |
| 47 | Vannest et al. (2012) | 15 | 6/9 | 15/0 | 9.2 (NA) | 7–14 | Animal judgement vs. tone detection | Knowledge | 4 |
| 48 | Wong et al. (2019) | 38 | 0/38 | 38/0 | 11.9 (1.0) | NA | Word relatedness judgement (multiple contrasts) | Relatedness | 18 |
| 49 | Wu et al. (2016) | 30 | 20/10 | 30/0 | 5.5 (0.3) | 5–5 | Sentence listening (prototypical, neutral, non-prototypical) | Knowledge | 12 |
| 50 | Xue et al. (2004) | 12 | 6/6 | 12/0 | 11.6 (NA) | 10–12 | Word relatedness judgment vs. fixation | Relatedness | 18 |

*Note.* Multiple experiments were derived from the same original article if and only if they were testing independent groups of children (see Section 2.1 and Turkeltaub et al., 2012). Additional information about the experiments can be found in the Appendix. # = experiment ID, *n* = sample size, F = female, M = male, R = right-handed, L = left-handed, no. = number, NA = not available.

### 3.2 Activation Likelihood Estimation

For general semantic cognition in children, the meta-analysis using ALE revealed spatial convergence of activation across experiments in eight clusters distributed across different regions of children’s cortex (see Table 2 and Figure 3). Ordered by cluster size, they were located in the left inferior frontal and precentral gyri (Clusters #1 and #8), the bilateral supplementary motor areas (Cluster #2), the left fusiform gyrus (Cluster #3), the right insular and inferior frontal cortices (Cluster #4), the left middle and superior temporal cortices (Cluster #5), the right inferior occipital gyrus and calcarine sulcus (Cluster #6), and the right fusiform gyrus (Cluster #7).

**Figure 3.**
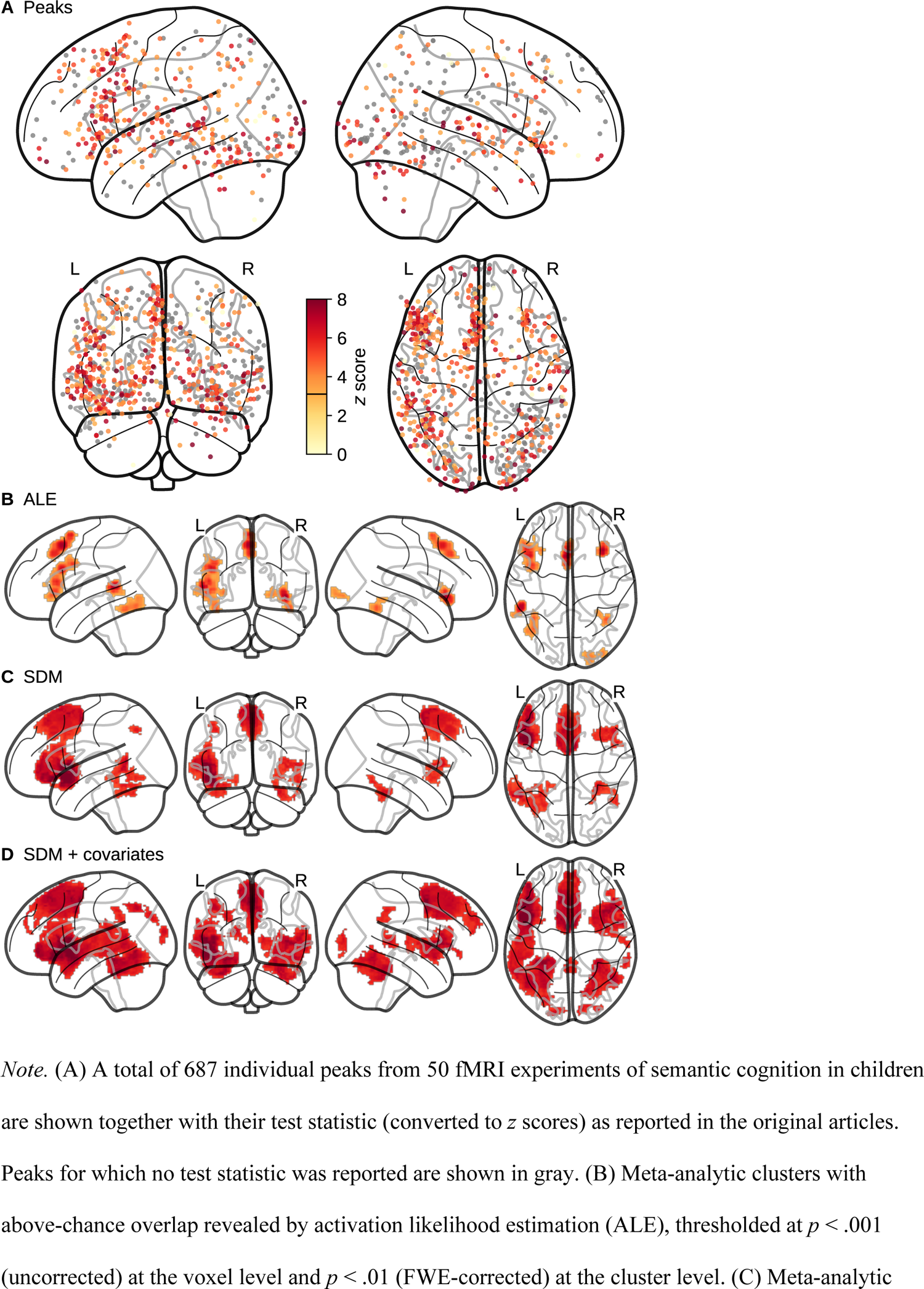
Meta-Analytic Results for Semantic Cognition in Children

**Table 2.**
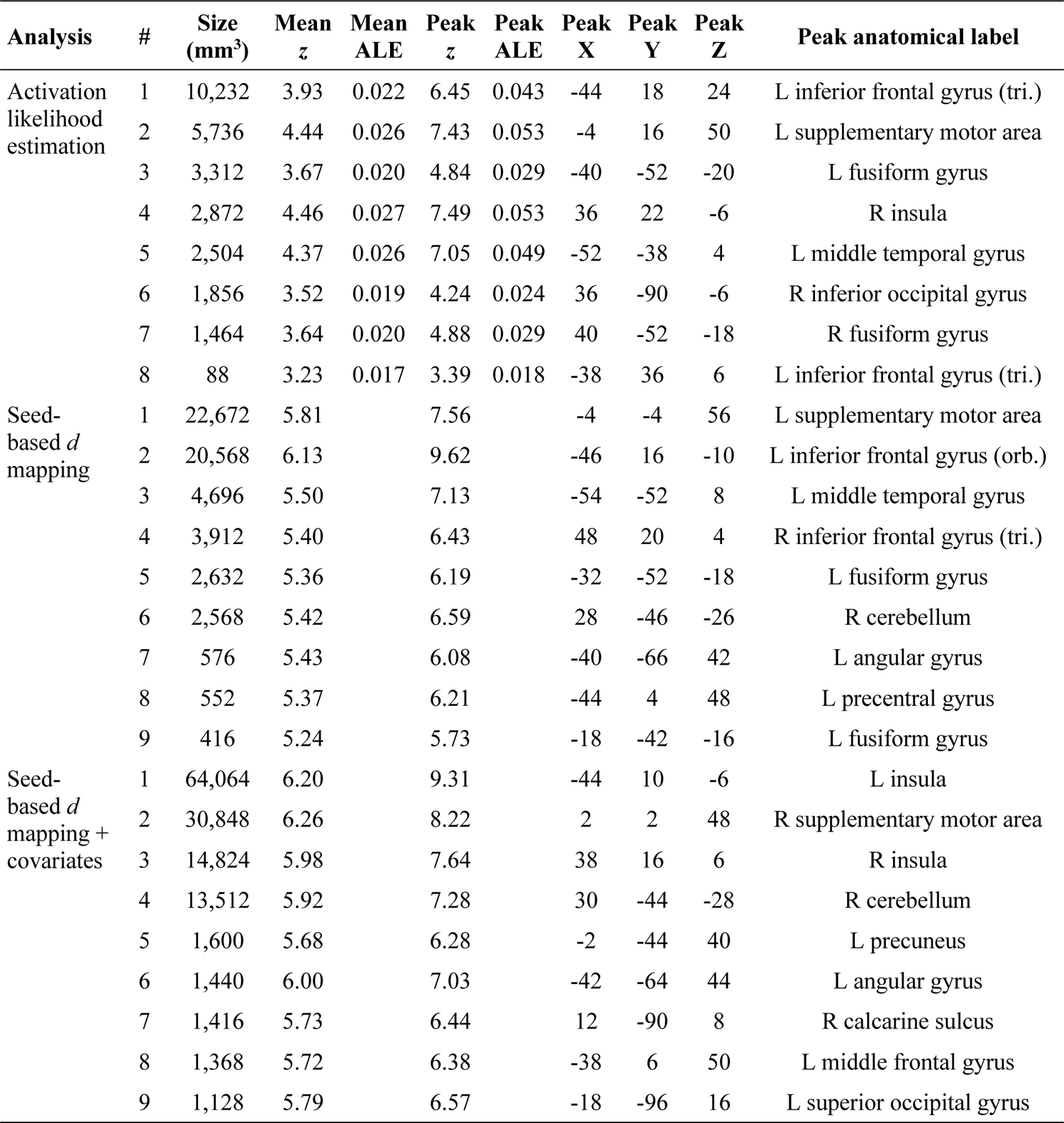
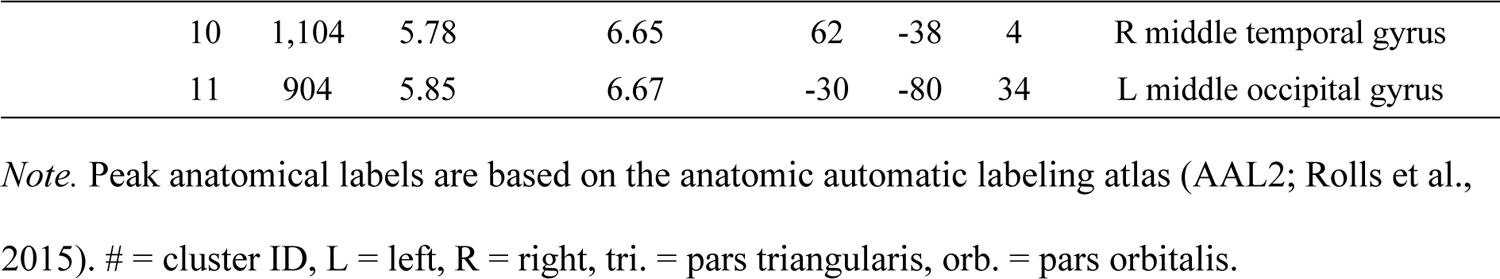
Statistics of the Meta-Analytic Clusters Shown in *Figure 3*

| Analysis | # | Size<br>($\text{mm}^3$) | Mean<br>$z$ | Mean<br>ALE | Peak<br>$z$ | Peak<br>ALE | Peak<br>X | Peak<br>Y | Peak<br>Z | Peak anatomical label |
| --- | --- | --- | --- | --- | --- | --- | --- | --- | --- | --- |
| Activation<br>likelihood<br>estimation | 1 | 10,232 | 3.93 | 0.022 | 6.45 | 0.043 | -44 | 18 | 24 | L inferior frontal gyrus (tri.) |
|  | 2 | 5,736 | 4.44 | 0.026 | 7.43 | 0.053 | -4 | 16 | 50 | L supplementary motor area |
|  | 3 | 3,312 | 3.67 | 0.020 | 4.84 | 0.029 | -40 | -52 | -20 | L fusiform gyrus |
|  | 4 | 2,872 | 4.46 | 0.027 | 7.49 | 0.053 | 36 | 22 | -6 | R insula |
|  | 5 | 2,504 | 4.37 | 0.026 | 7.05 | 0.049 | -52 | -38 | 4 | L middle temporal gyrus |
|  | 6 | 1,856 | 3.52 | 0.019 | 4.24 | 0.024 | 36 | -90 | -6 | R inferior occipital gyrus |
|  | 7 | 1,464 | 3.64 | 0.020 | 4.88 | 0.029 | 40 | -52 | -18 | R fusiform gyrus |
|  | 8 | 88 | 3.23 | 0.017 | 3.39 | 0.018 | -38 | 36 | 6 | L inferior frontal gyrus (tri.) |
| Seed-based $d$<br>mapping | 1 | 22,672 | 5.81 | | 7.56 | | -4 | -4 | 56 | L supplementary motor area |
|  | 2 | 20,568 | 6.13 |  | 9.62 |  | -46 | 16 | -10 | L inferior frontal gyrus (orb.) |
|  | 3 | 4,696 | 5.50 |  | 7.13 |  | -54 | -52 | 8 | L middle temporal gyrus |
|  | 4 | 3,912 | 5.40 |  | 6.43 |  | 48 | 20 | 4 | R inferior frontal gyrus (tri.) |
|  | 5 | 2,632 | 5.36 |  | 6.19 |  | -32 | -52 | -18 | L fusiform gyrus |
|  | 6 | 2,568 | 5.42 |  | 6.59 |  | 28 | -46 | -26 | R cerebellum |
|  | 7 | 576 | 5.43 |  | 6.08 |  | -40 | -66 | 42 | L angular gyrus |
|  | 8 | 552 | 5.37 |  | 6.21 |  | -44 | 4 | 48 | L precentral gyrus |
|  | 9 | 416 | 5.24 |  | 5.73 |  | -18 | -42 | -16 | L fusiform gyrus |
| Seed-based $d$<br>mapping +<br>covariates | 1 | 64,064 | 6.20 | | 9.31 | | -44 | 10 | -6 | L insula |
|  | 2 | 30,848 | 6.26 |  | 8.22 |  | 2 | 2 | 48 | R supplementary motor area |
|  | 3 | 14,824 | 5.98 |  | 7.64 |  | 38 | 16 | 6 | R insula |
|  | 4 | 13,512 | 5.92 |  | 7.28 |  | 30 | -44 | -28 | R cerebellum |
|  | 5 | 1,600 | 5.68 |  | 6.28 |  | -2 | -44 | 40 | L precuneus |
|  | 6 | 1,440 | 6.00 |  | 7.03 |  | -42 | -64 | 44 | L angular gyrus |
|  | 7 | 1,416 | 5.73 |  | 6.44 |  | 12 | -90 | 8 | R calcarine sulcus |
|  | 8 | 1,368 | 5.72 |  | 6.38 |  | -38 | 6 | 50 | L middle frontal gyrus |
|  | 9 | 1,128 | 5.79 |  | 6.57 |  | -18 | -96 | 16 | L superior occipital gyrus |

|  |  |  |  |  |  |  |  |
| --- | --- | --- | --- | --- | --- | --- | --- |
| 10 | 1,104 | 5.78 | 6.65 | 62 | -38 | 4 | R middle temporal gyrus |
| 11 | 904 | 5.85 | 6.67 | -30 | -80 | 34 | L middle occipital gyrus |

*Note.* Peak anatomical labels are based on the anatomic automatic labeling atlas (AAL2; Rolls et al., 2015). # = cluster ID, L = left, R = right, tri. = pars triangularis, orb. = pars orbitalis.

### 3.3 Seed-Based *d* Mapping

Meta-analytic effect size maps were created based on the test statistics (*z* scores or *t* scores) of the reported peak coordinates. These were available for 461 (67.1%) of all peak coordinates, whereas the test statistics for the remaining 226 peak coordinates were inferred via multiple imputations as described in Albajes-Eizagirre et al. (2019). This alternative meta-analytic approach yielded qualitatively similar results as ALE: The largest clusters (and highest effect sizes) were again observed in the left inferior frontal gyrus, the bilateral supplementary motor areas, and the left middle and superior temporal gyri (see Figure 3C and Table 2). Three noteworthy differences between the results from these two different meta-analytic algorithms were (a) that the size of the significant clusters was larger overall for SDM as compared to ALE, (b) that one cluster in the right visual cortex was observed with ALE but not with SDM, and (c) that one cluster in the left angular gyrus was observed with SDM but not with ALE.

The effect size-based approach not only served as a robustness check for the main (ALE) analysis but also made it possible to re-assess the results while controlling for four different linear covariates of no interest (namely the mean age of the sample, the modality of stimulus presentation, the modality of children’s response, and the statistical software package used for data analysis). This again yielded qualitatively similar results, although cluster sizes were larger than in the original analysis without covariates (see Figure 3D and Table 2).

### 3.4 Differences Between Semantic Task Categories

Task category-specific sub-analyses for experiments probing semantic world knowledge (e.g., naming a word after hearing its description) and for experiments probing semantic relatedness (e.g., hearing two words and deciding if they are related or not) both showed the largest clusters of activation in the bilateral supplementary motor areas, in the pars triangularis of the left inferior frontal gyrus, and in the right insular and inferior frontal cortices (see Figure 4 and Table 3). For experiments probing semantic relatedness, there was one additional cluster in the left middle temporal gyrus. The sub-analysis for experiments probing the discrimination of visual semantic object categories showed three clusters of consistent activation in the bilateral fusiform gyri as well as in the visual cortex of the right occipital lobe.

**Table 3.**
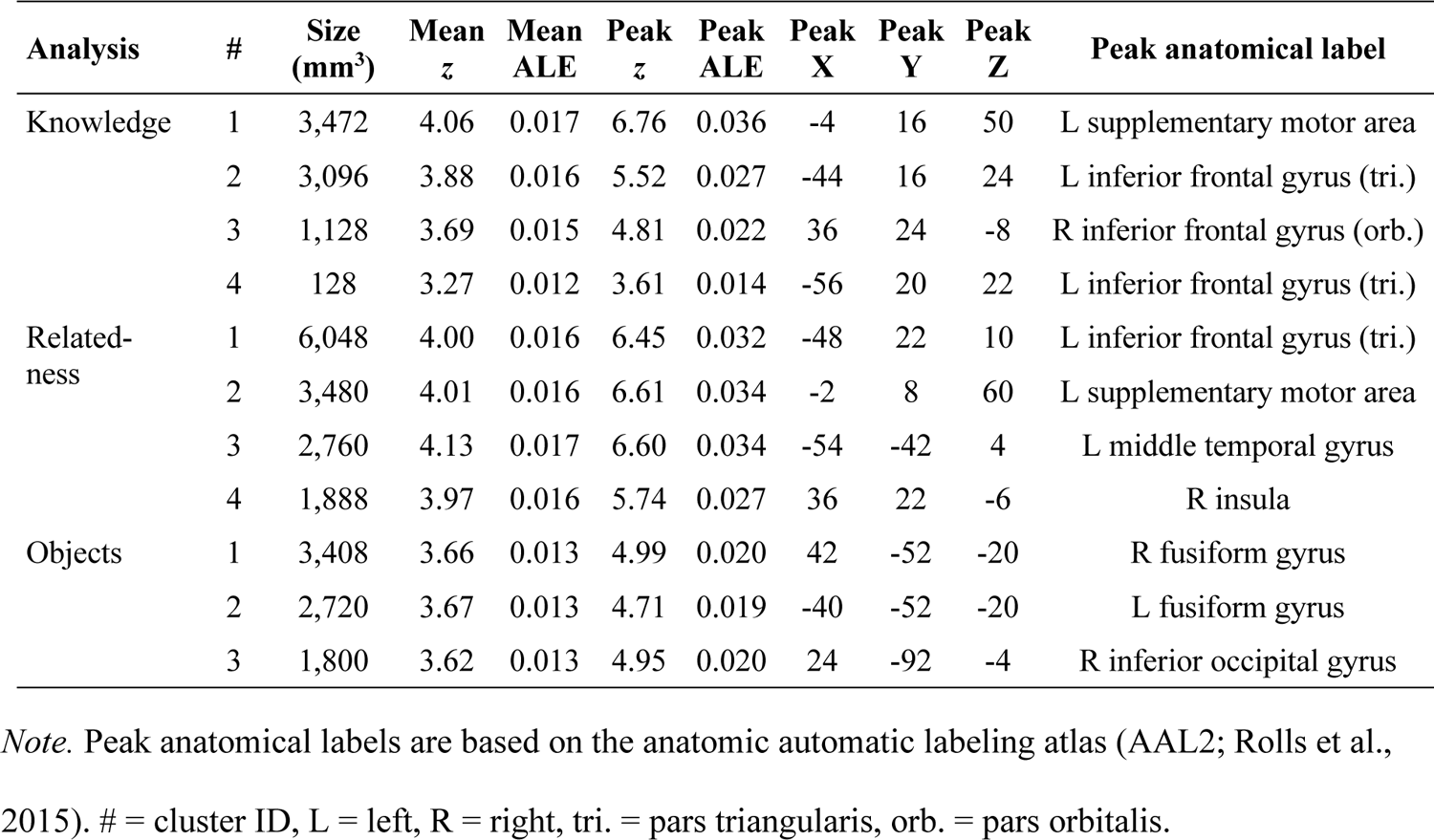
Statistics for the Meta-Analytic Clusters Shown in *Figure 4*

**Figure 4.**
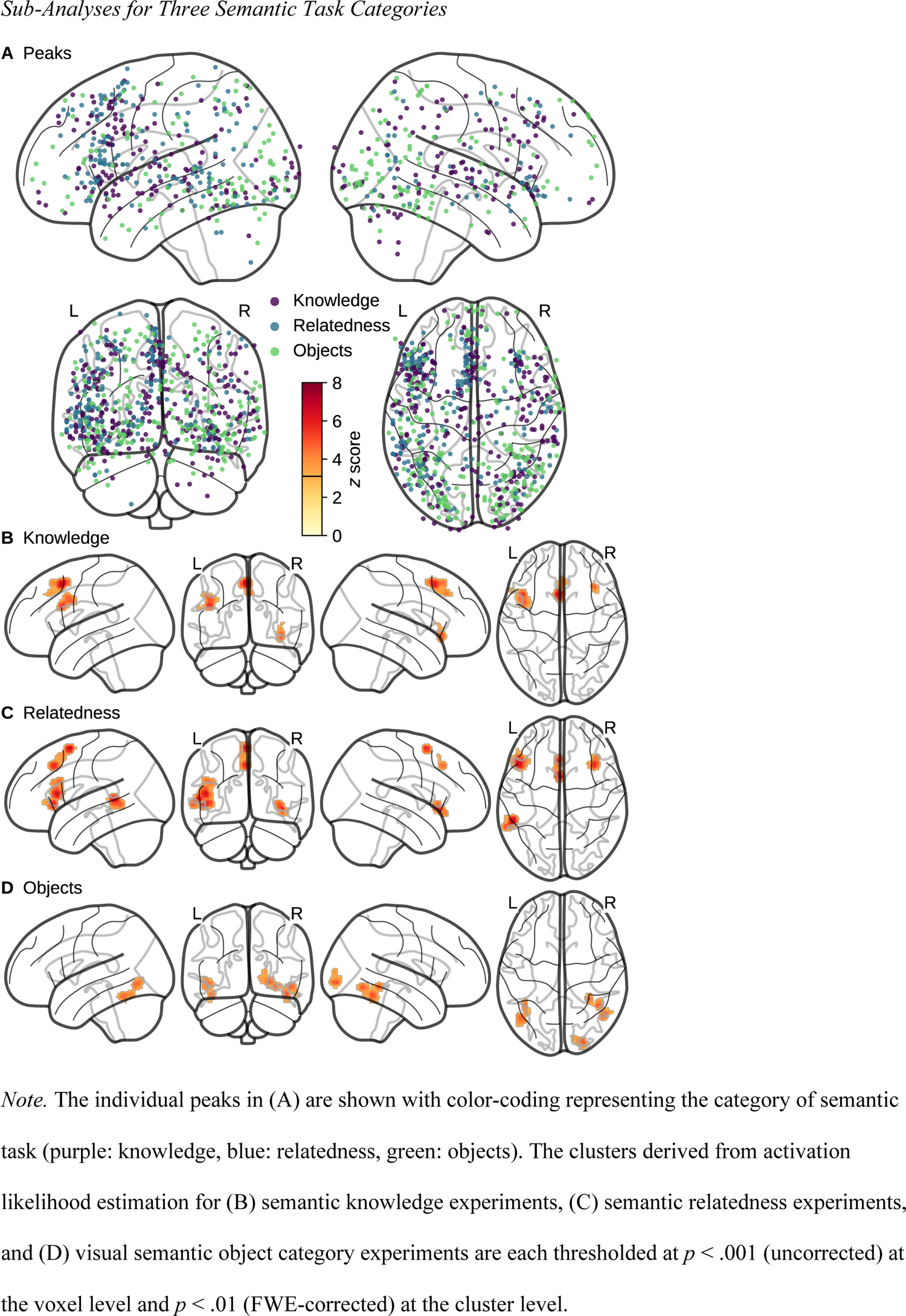
Sub-Analyses for Three Semantic Task Categories

These task-specific meta-analytic maps were contrasted against one another to examine where they differed reliably from one another (see Figure 5 and Table 4). First, experiments probing semantic knowledge showed more consistent activation than the other two task categories in two small clusters in the left insular and middle frontal cortices. Second, tasks probing semantic relatedness showed more consistent activation than the other two task categories in the pars opercularis of the left inferior frontal gyrus and in the left middle temporal gyrus. Finally, tasks probing visual semantic object categories showed more consistent activation than the other two task categories in the bilateral occipital and fusiform cortices as well as in the right superior parietal cortex. They also showed reliably less activation than the other two task categories in one small cluster in the left medial frontal lobe (shown in blue in Figure 5C). The conjunction analysis revealed that there were no regions of activation that were shared by all three semantic task categories (as seen also in Figure 4B–D).

**Figure 5.**
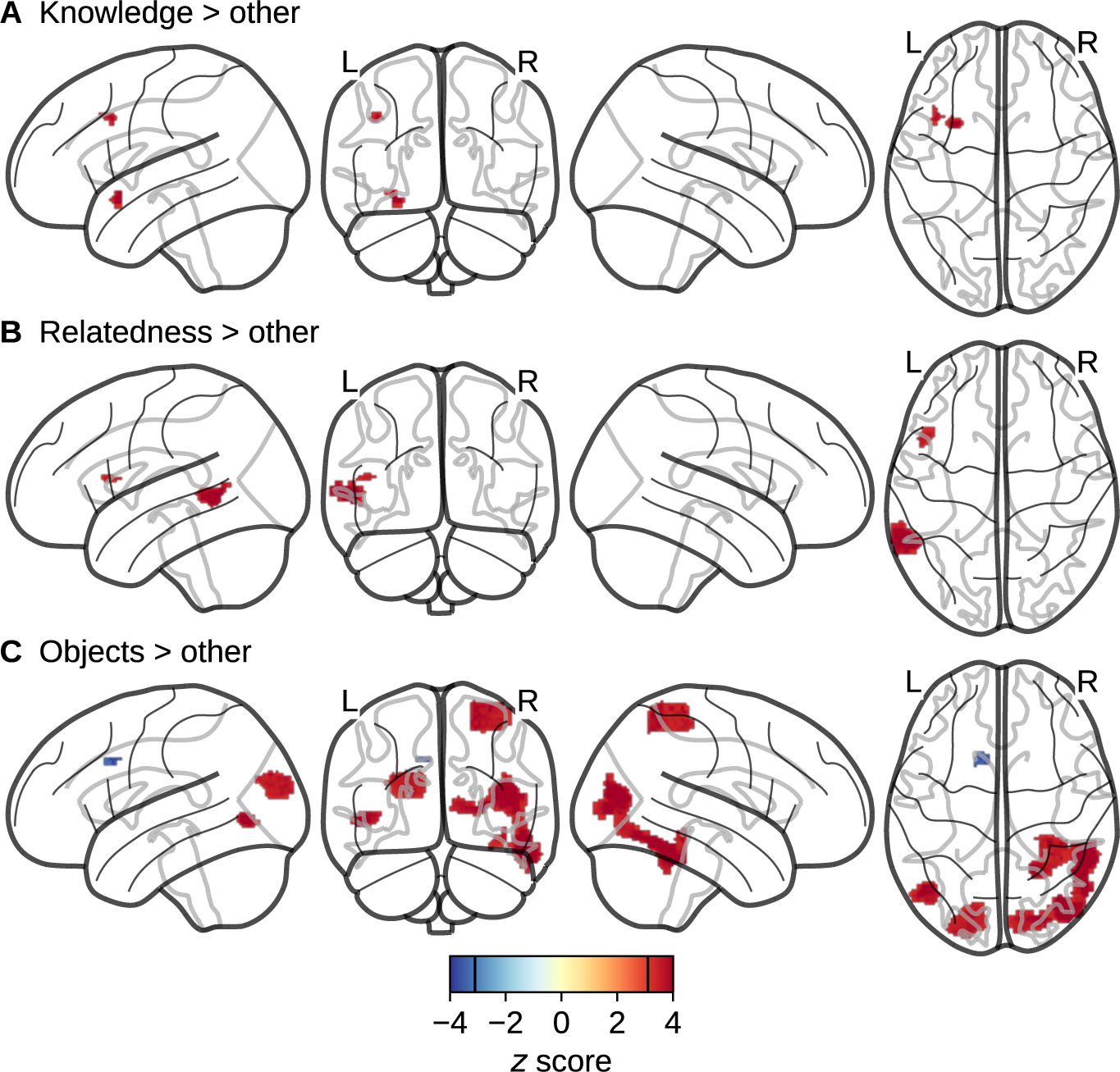
Differences Between Semantic Task Categories

**Table 4.** Statistics for the Meta-Analytic Clusters Shown in *Figure 5*

| Analysis | # | Size<br>($\text{mm}^3$) | Mean<br>$z$ | Peak<br>$z$ | Peak<br>X | Peak<br>Y | Peak<br>Z | Peak anatomical label |
| --- | --- | --- | --- | --- | --- | --- | --- | --- |
| Knowledge ><br>(relatedness + objects) | 1 | 304 | 3.62 | 4.06 | -28 | 10 | -16 | L insula |
|  | 2 | 256 | 3.49 | 3.89 | -40 | 14 | 36 | L middle frontal gyrus |
| Relatedness ><br>(knowledge + objects) | 1 | 1,960 | 3.72 | 4.06 | -58 | -50 | 0 | L middle temporal gyrus |
|  | 2 | 336 | 3.42 | 3.62 | -46 | 14 | 12 | L inferior frontal gyrus (oper.) |
| Objects > (knowledge<br>+ relatedness) | 1 | 9,912 | 3.71 | 4.06 | 46 | -68 | -4 | R middle inferior gyrus |
|  | 2 | 4,744 | 3.48 | 4.06 | 28 | -46 | 66 | R postcentral gyrus |
|  | 3 | 4,048 | 3.45 | 3.89 | -18 | -84 | 26 | L superior occipital gyrus |
|  | 4 | 1,104 | 3.65 | 4.06 | -46 | -70 | -2 | L middle occipital gyrus |
|  | 5 | 312 | 3.48 | 4.06 | 38 | -40 | -14 | R fusiform gyrus |
|  | 6 | 200 | -3.52 | -3.54 | -10 | 12 | 34 | L middle cingulate gyrus |
| Conjunction | *No significant clusters* |  |  |  |  |  |  |  |

*Note.* Peak anatomical labels are based on the anatomic automatic labeling atlas (AAL2; Rolls et al., 2015). # = cluster ID, L = left, R = right, oper. = pars opercularis.

### 3.5 Age-Related Changes

Age-related changes in fMRI activation patterns across the 50 experiments were examined (a) by splitting the sample of experiments in half at the median of the (mean) sample ages (*md*_mean age_ = 10.3 years) and performing an ALE subtraction analysis as described above (see Section 2.4) and (b) by entering mean sample age as a linear predictor in a meta-regression model using the effect size-based SDM approach (see Section 2.3).

The median split-based approach showed more consistent activation in experiments with older (> 10.3 years) as compared to younger (< 10.3 years) children only at the right putamen and insula (see Figure 6A and Table 5). The effect size-based approach showed no age-related changes using the pre-specified statistical threshold (*p* < .001 [FWE-corrected] at the voxel level and *k* > 25 connected voxels [200 mm^3^] at the cluster level).

**Figure 6.**
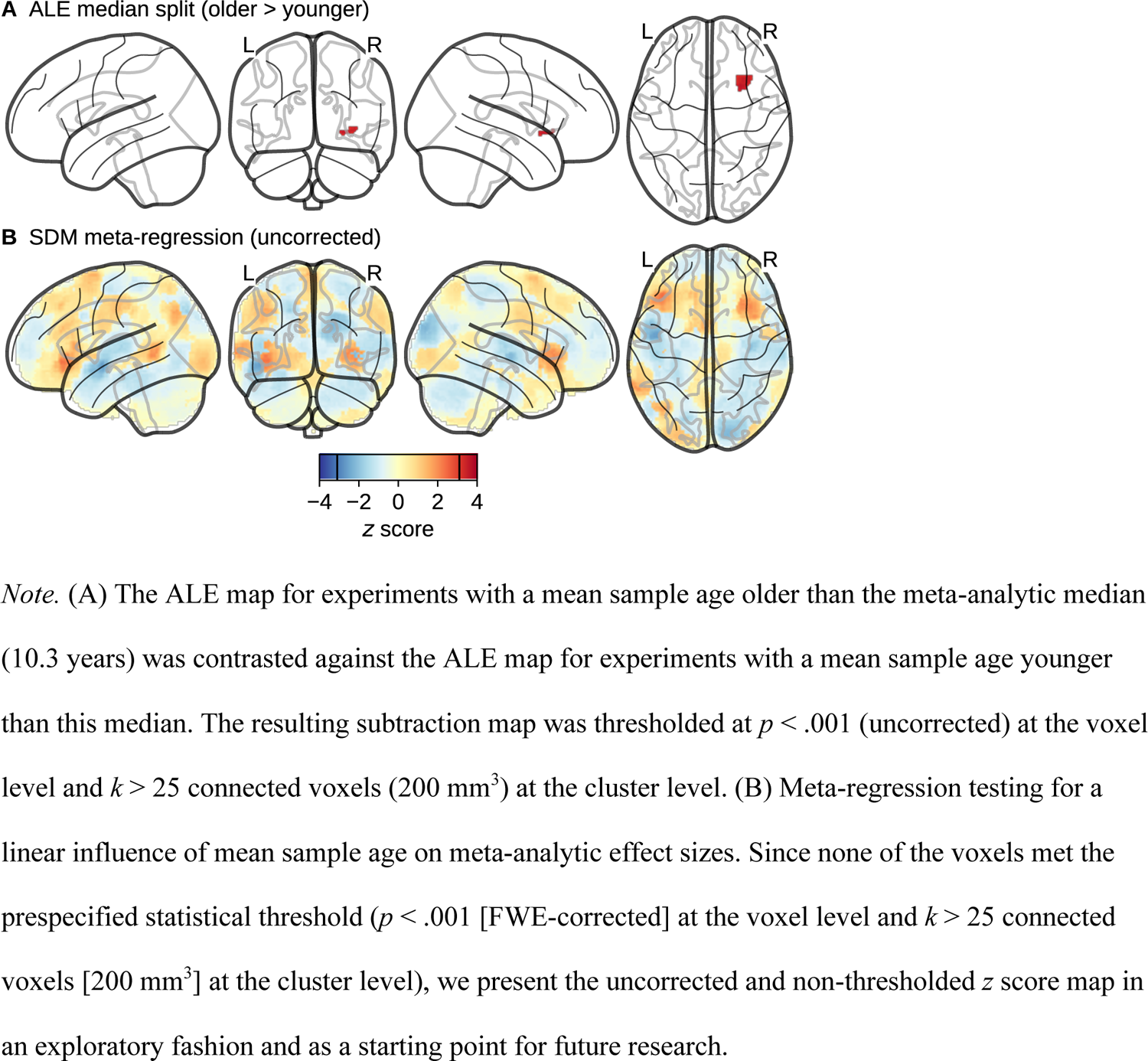
Differences Between Older and Younger Children

**Table 5.** Statistics for the Meta-Analytic Clusters Shown in *Figure 6*

Statistics for the Meta-Analytic Clusters Shown in Figure 6
| Analysis | # | Size<br>(mm<sup>3</sup>) | Mean<br>z | Peak<br>z | Peak<br>X | Peak<br>Y | Peak<br>Z | Peak anatomical label |
| --- | --- | --- | --- | --- | --- | --- | --- | --- |
| Median split (older > younger) | 1 | 536 | 3.41 | 3.62 | 28 | 12 | -6 | R putamen |
| Meta-regression |  |  |  |  | No significant clusters |  |  |  |

*Note.* Peak anatomical labels are based on the anatomic automatic labeling atlas (AAL2; Rolls et al., 2015). # = cluster ID, R = right.

However, one should consider that meta-analytic null effects for study-level moderating variables may in part reflect the lack of statistical power for detecting them (Hempel et al., 2013). This is especially true when the variable of interest (here: mean sample age) has a restricted variance (see Figure 2 and Table 1). To mitigate this lack of statistical power in a post hoc fashion, we present the uncorrected and non-thresholded *z* score map from the effect size-based meta-regression in Figure 6B. This map suggests an age-related decrease of effect sizes in the left middle/superior temporal gyrus (peak *z* = −2.55) and an age-related increase of effect sizes in the left inferior frontal gyrus (peak *z* = 2.33). To a lesser extent, the increase in the inferior frontal gyrus is mirrored in the right hemisphere, consistent with the median split-based result from ALE. However, none of these peaks survived our initial cluster-forming threshold and therefore additional experiments will be needed to confirm if these age-related changes turn out to be reliable on a meta-analytic level.

### 3.6 Comparison With Semantic Cognition in Adults

A recent meta-analysis by Jackson (2021) used ALE to synthesize the fMRI literature on semantic control in adults. They also broadened their analysis to 415 studies of general semantic cognition and found wide-ranging clusters of consistent activation especially in the left hemisphere, spanning multiple areas in the temporal and inferior frontal lobes as well as the supplementary motor area (see Figure 7A and Table 6 for a reproduction of these results based on the peak coordinates kindly provided by the original author). This meta-analytic map of semantic cognition in adults was compared to the map of semantic cognition in children by means of a subtraction and conjunction analysis. This revealed more consistent activation in children as compared to adults in multiple posterior regions of the cortex, including the bilateral inferior temporal and right inferior parietal gyri (see Figure 7B and Table 6). In contrast, adults showed more consistent activation in the anterior part of the left middle and inferior temporal gyri as well as deep in the left calcarine sulcus. Finally, the conjunction analysis indicated large areas of overlap between the two groups in the left inferior frontal gyrus, the supplementary motor area, the left middle and superior temporal gyri, the right insular and inferior frontal cortices, and the left fusiform gyrus (see Figure 7C and Table 6).

**Figure 7.**
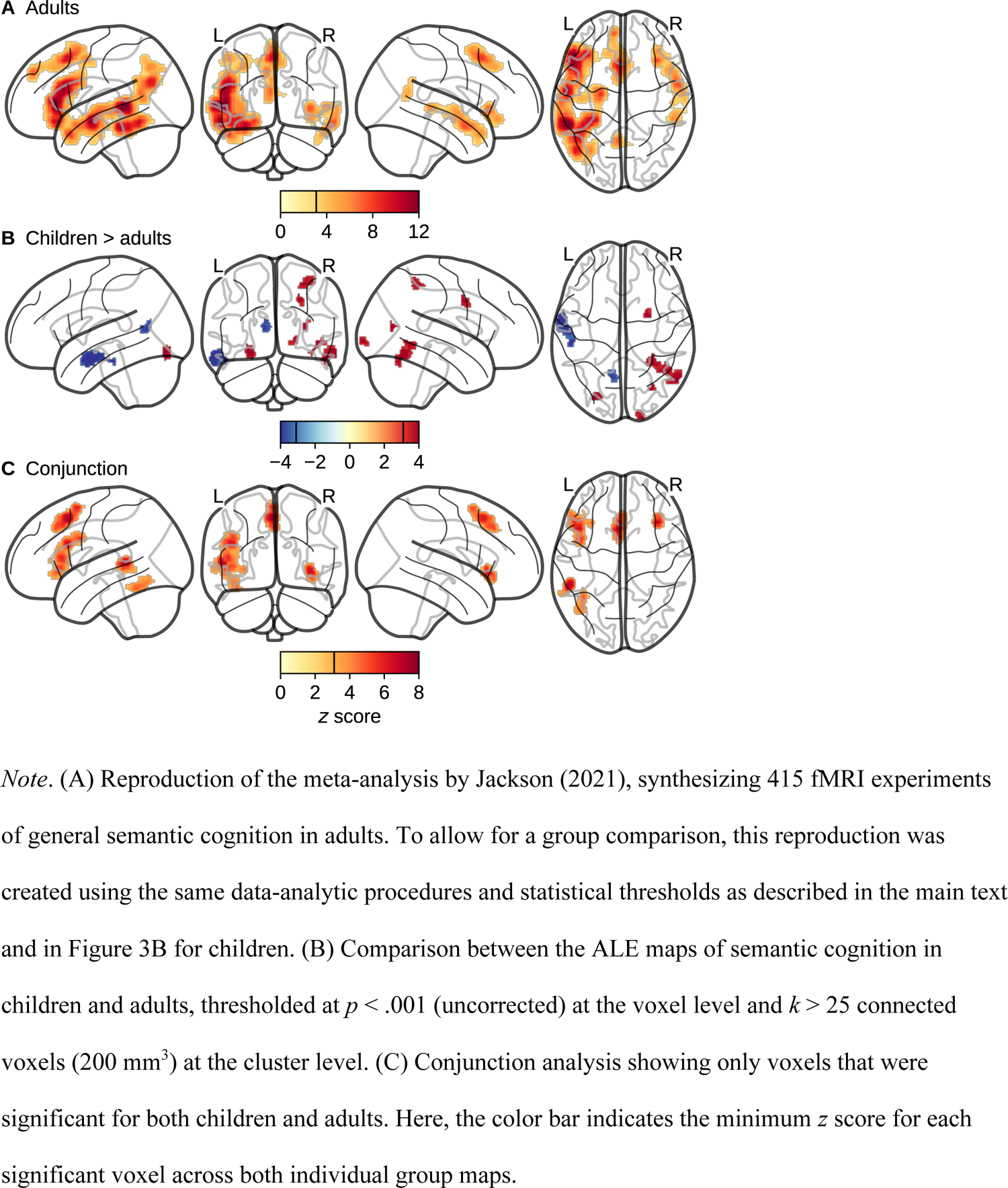
Comparison With Semantic Cognition in Adults

**Table 6.** Statistics for the Meta-Analytic Clusters Shown in *Figure 7*

| Analysis | # | Size<br>(mm<sup>3</sup>) | Mean<br>z | Mean<br>ALE | Peak<br>z | Peak<br>ALE | Peak<br>X | Peak<br>Y | Peak<br>Z | Peak anatomical label |
| --- | --- | --- | --- | --- | --- | --- | --- | --- | --- | --- |
| Adults | 1 | 88,648 | 5.41 | 0.064 | 13.27 | 0.179 | -56 | -38 | 2 | L middle temporal gyrus |
|  | 2 | 12,488 | 4.65 | 0.054 | 9.68 | 0.121 | -4 | 18 | 50 | L supplementary motor area |
|  | 3 | 8,320 | 4.35 | 0.050 | 7.54 | 0.090 | 56 | 0 | -18 | R middle temporal gyrus |
|  | 4 | 4,176 | 4.35 | 0.050 | 7.37 | 0.087 | 36 | 24 | -2 | R insula |
|  | 5 | 3,616 | 4.36 | 0.050 | 6.89 | 0.081 | -6 | -56 | 14 | L precuneus |
|  | 6 | 1,768 | 3.99 | 0.046 | 6.11 | 0.071 | 52 | -34 | 0 | R middle temporal gyrus |
|  | 7 | 72 | 3.22 | 0.038 | 3.35 | 0.039 | -26 | 26 | 46 | L middle frontal gyrus |
| Children<br>> adults | 1 | 1,072 | 3.68 |  | 3.89 |  | 56 | -64 | -18 | R inferior temporal gyrus |
|  | 2 | 776 | 3.59 |  | 3.89 |  | 30 | -48 | 52 | R inferior parietal gyrus |
|  | 3 | 672 | 3.59 |  | 3.48 |  | 44 | -56 | -8 | R inferior temporal gyrus |
|  | 4 | 664 | 3.69 |  | 4.06 |  | -26 | -82 | -16 | L lingual gyrus |
|  | 5 | 432 | 3.51 |  | 4.06 |  | 20 | -104 | -2 | R calcarine sulcus |
|  | 6 | 352 | 3.66 |  | 4.06 |  | 26 | 0 | 36 | No label found |
|  | 7 | 232 | 3.69 |  | 3.39 |  | 32 | -74 | 10 | R middle occipital gyrus |
| Adults<br>> children | 1 | 2,632 | 3.78 |  | 4.06 |  | -58 | -10 | -22 | L middle temporal gyrus |
|  | 2 | 584 | 3.49 |  | 4.06 |  | -10 | -62 | 10 | L calcarine sulcus |
| Conjunction | 1 | 9,592 | 3.95 | 0.023 | 6.45 | 0.043 | -44 | 18 | 24 | L inferior frontal gyrus (tri.) |
|  | 2 | 5,320 | 4.41 | 0.027 | 7.43 | 0.053 | -4 | 16 | 50 | L supplementary motor area |
|  | 3 | 2,480 | 4.38 | 0.026 | 7.05 | 0.049 | -52 | -38 | 4 | L middle temporal gyrus |
|  | 4 | 2,232 | 4.23 | 0.028 | 6.74 | 0.053 | 36 | 24 | -4 | R insula |
|  | 5 | 2,056 | 3.66 | 0.021 | 4.84 | 0.029 | -40 | -52 | -20 | L fusiform gyrus |
|  | 6 | 48 | 3.20 | 0.017 | 3.30 | 0.018 | -40 | 36 | 6 | L inferior frontal gyrus (tri.) |

*Note.* Peak anatomical labels are based on the anatomic automatic labeling atlas (AAL2; Rolls et al., 2015). # = cluster ID, L = left, R = right, tri. = pars triangularis.

### 3.7 Evaluation of Robustness

The robustness of the meta-analytic results against two different types of publication bias—spurious findings and the file drawer problem—was assessed using a leave-one-out (jackknife) analysis and a fail-safe N analysis. Both of these analyses were conducted for the entire sample of all 50 semantic experiments (see Figure 3B and Table 2) and for each of the three task category-specific sub-analyses (see Figure 4B–D and Table 3).

The leave-one-out procedure showed that all clusters detected in the main analysis were robust against the deletion of individual studies, with an average leave-one-out robustness of 96% across the eight clusters (range 84–100%; see Figure 8). For the sub-analysis of semantic relatedness experiments, the robustness of all four clusters was at 100%, whereas for semantic knowledge experiments, it was at 100% for Clusters #1 and #2 but reduced for Clusters #3 (right insula; 48%) and #4 (left IFG, 81%). Finally, for visual semantic object category experiments, it was slightly reduced (85%) for all three clusters. Together, this reflects good overall robustness against spurious experiments in the meta-analysis, although this robustness was compromised slightly for the sub-analyses that were run on fewer experiments (Eickhoff et al., 2016).

**Figure 8.**
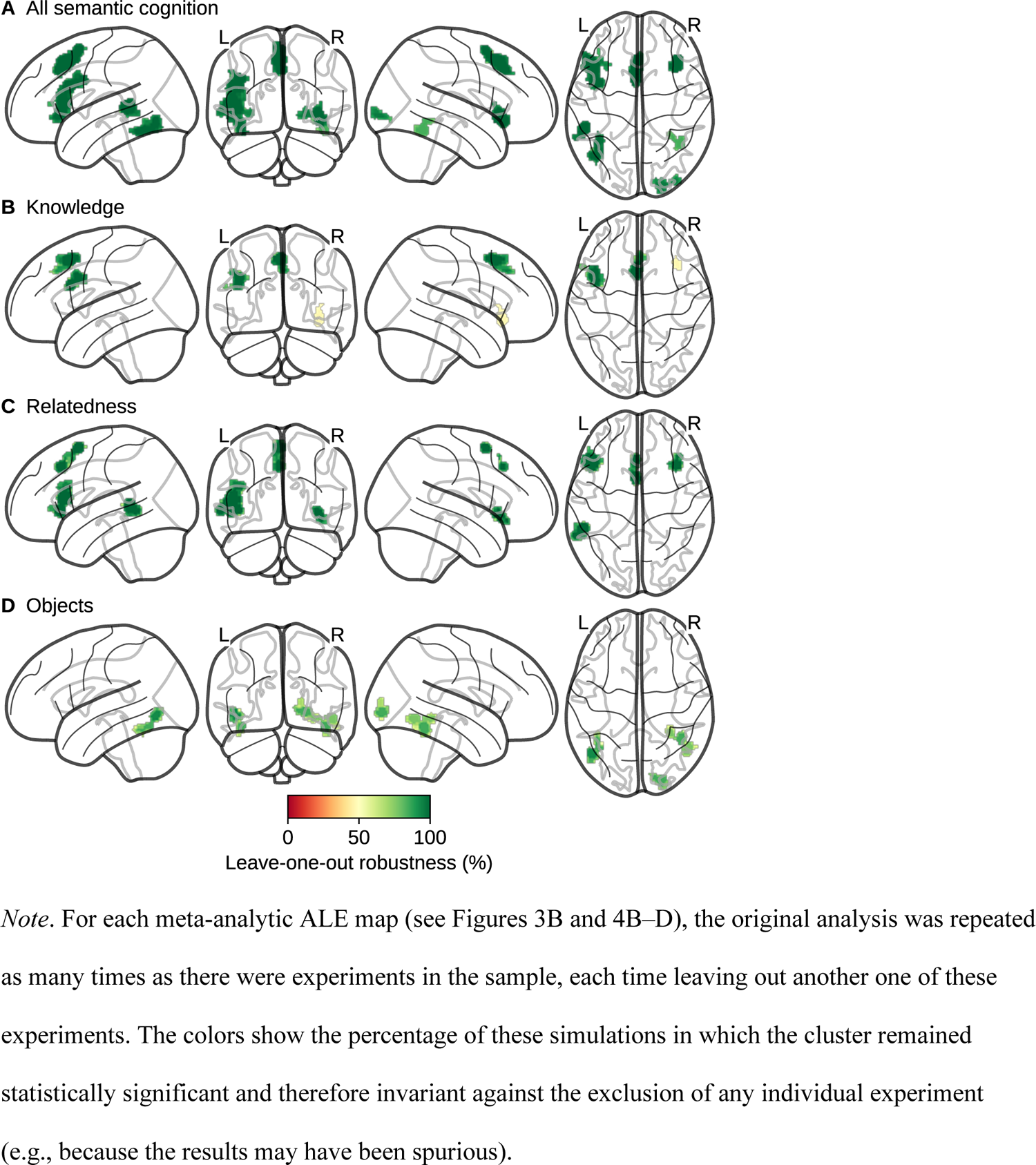
Leave-One-Out Analysis

The fail-safe N analysis showed that most clusters were robust against the file drawer problem. This was indicated by the fact that in these cases, the number of (unpublished) null experiments that needed to be added until overturning the statistical significance of the cluster exceeded the number of (published) experiments in the original analysis (see Figure 9). The only clusters were this was not the case were Clusters #5 (right occipital lobe; FSN = 18), #6 (right fusiform gyrus; FSN = 15), and #8 (left inferior frontal gyrus; FSN = 1). Note, however, that Clusters #5 and #6 still marginally exceeded the desired value of 30% of the original sample size (see Section 2.7 and Samartsidis et al., 2020). For the task category-specific sub-analyses, FSN values were high overall, except for the knowledge-related Clusters #3 (right insula; FSN = 1) and #4 (left inferior frontal gyrus; FSN = 2) as well as the object-related Clusters #1 (right fusiform gyrus; FSN = 8), #2 (left fusiform gyrus; FSN = 5), and #3 (right occipital lobe; FSN = 11). All but the two knowledge-related clusters exceeded the desired threshold of 30%, once more giving the overall impression of satisfactory robustness to publication bias. However, these two specific clusters need to be interpreted with caution and would require additional support by future fMRI experiments.

**Figure 9.**
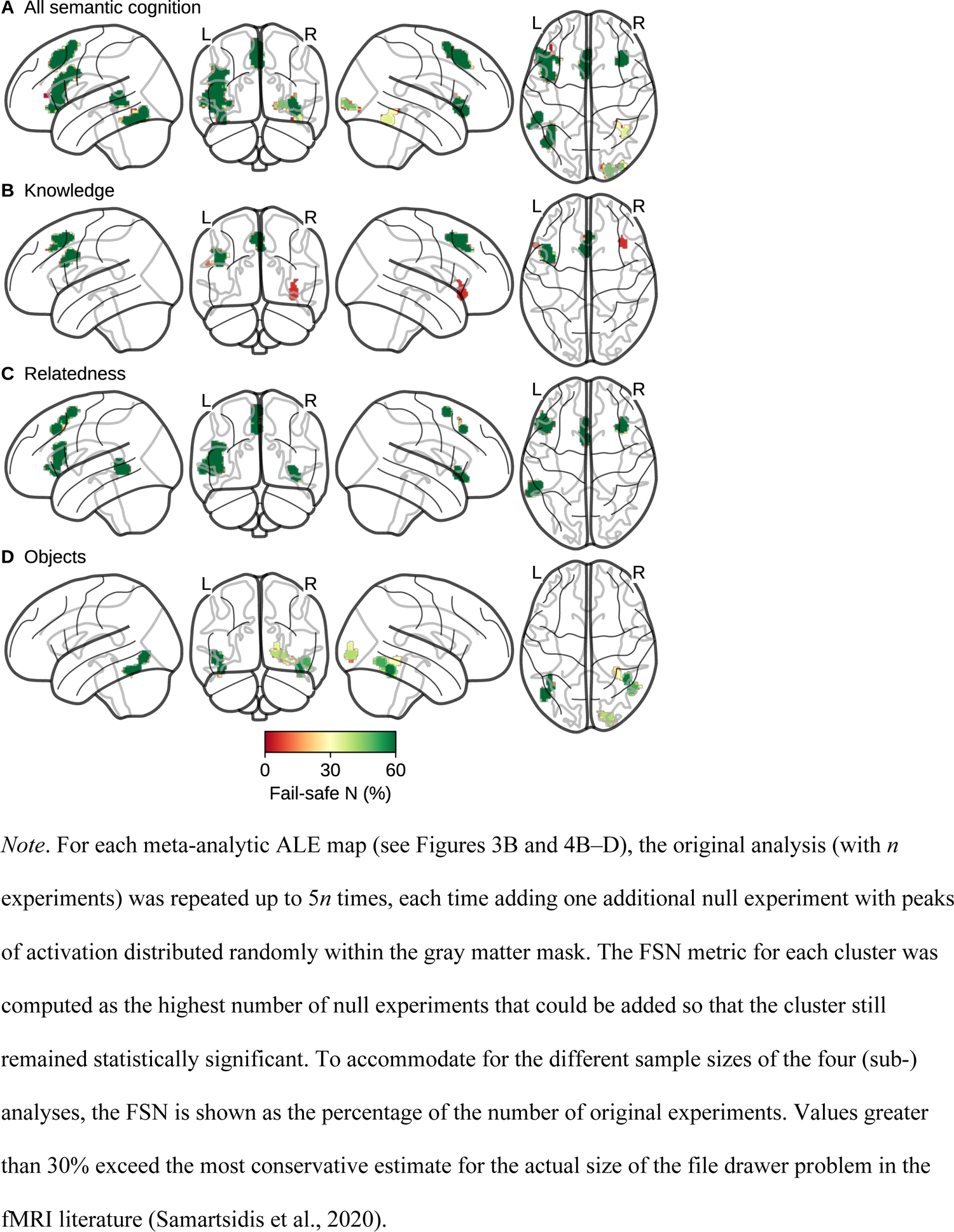
Fail-Safe N Analysis

## 4. Discussion

Here we systematically localized the brain areas underlying semantic cognition in children by means of a coordinate-based meta-analysis of fMRI studies. We identified 50 individual experiments scanning children with a mean age of 3–12 years using a variety of semantic tasks. Pooling across the reported peak coordinates from all of these experiments, we found evidence for consistent activation in sub-regions of the left perisylvian language network associated with lexical processing (left MTG/STG and IFG) as well as in the bilateral SMA, the right insula, and more posterior brain regions in the bilateral fusiform and right occipital cortices. These areas were recruited to a different degree by different semantic task categories: Inferior frontal regions and the SMA were recruited preferentially during tasks tapping into semantic knowledge (e.g., naming an object after hearing its descriptions) and semantic relatedness (e.g., hearing two words and deciding if they are related or not), while posterior regions were recruited preferentially during tasks tapping into the differentiation of visual object categories (e.g., passively viewing faces as compared to other visual stimuli).

The left MTG/STG and the pars triangularis of the left IFG are known to be implicated in semantic processing from at least 3 years of age onwards (Skeide et al., 2014). There is also some evidence that children are able to process word meaning in the left MTG with as little as 2 years of age (Friedrich & Friederici, 2010; Travis et al., 2011). Within the left IFG, semantic processing especially recruits the more anterior parts (pars triangularis and pars orbitalis; Brauer & Friederici, 2007; Nuñez et al., 2011; Skeide et al., 2014; Skeide & Friederici, 2016) which also showed the strongest meta-analytic peaks in our study. These sub-areas seem to play an especially crucial role in children’s language processing, as they allow them to successfully retrieve the semantic meaning of grammatically challenging sentences even though their syntactic abilities (localized in the pars opercularis of the left IFG in adults) are not yet fully developed (Skeide et al., 2014).

Just as the left IFG and MTG/STG, the bilateral SMA also showed meta-analytically robust activation in children performing semantic knowledge and relatedness tasks. This mirrors previous meta-analyses of semantic cognition in adults (e.g., Binder et al., 2009; Jackson, 2021) as well as meta-analyses of language comprehension in both children (Enge et al., 2020; Martin et al., 2015) and adults (e.g., Ferstl et al., 2008; Rodd et al., 2015). The premotor activation could reflect a grounding of abstract semantic concepts in articulatory motor representations (Martin, 2016; Pulvermüller & Fadiga, 2010). Alternatively or in addition to this, activation in the anterior part of the SMA (pre-SMA) may also reflect higher-order cognitive control processes such as ambiguity resolution and the integration of semantic context (Hertrich et al., 2016). This is supported by our observation that the SMA showed no consistent activation when children performed visual object category tasks. These tasks would in many cases afford a similar degree of pre-motor response (e.g., when viewing tools; e.g., Dekker et al., 2014; Kersey et al., 2016) but arguably a lesser degree of cognitive control compared to tasks probing semantic knowledge or relatedness (see Binder et al., 2009, for similar findings focusing exclusively on experiments with linguistic stimuli in adults). Note, however, that only a limited number of visual object category tasks (*n* = 13) could be included in the present meta-analysis, presumably limiting statistical power (Eickhoff et al., 2016).

Finally, the ventral temporal cortex (fusiform gyrus and adjacent areas) is well-known to house category-selective neuronal populations that respond primarily to certain categories of visual stimuli (e.g., faces in the fusiform face area [FFA] or objects in the lateral occipital complex [LOC]; Grill-Spector & Weiner, 2014; Haxby et al., 2001). Accordingly, these regions showed consistent activation only for tasks in which children viewed different visual semantic object categories. A subset of small areas within this ventral temporal and occipital region also showed the most prominent increase of meta-analytic activation for children as compared to adults. This may reflect that children need to recruit these patches of cortex to a stronger degree than adults to distinguish between different kinds of visual stimuli (see also Antonucci & Alt, 2011). We cannot preclude, however, that this group difference could also be driven by differences in the kinds of tasks and baseline conditions chosen to investigate semantic cognition in children and adults. Furthermore, the semantic categorization of different kinds of visual objects is confounded with lower-level sensory differences between them. These visual confounds might at least partially explain our meta-analytic results for this task category (e.g., the activation of the right early visual cortex; see Figure 4D).

There was one region, namely the left ATL, that is oftentimes considered to be at the core of the semantic system (Lambon Ralph et al., 2017; Patterson et al., 2007) but did not show any meta-analytically consistent activation in children whatsoever. In adults, the ATL serves as an amodal “hub” connecting different modality-specific sites within the wider semantic network (e.g., speech processing in the IFG and visual semantics in the occipital and ventral temporal cortices).

Neuroimaging studies of semantic cognition in children seem to elicit significantly less of such ATL activation (see Figure 7B), suggesting that this semantic hub may need time to develop over childhood and into adolescence (Hwang et al., 2013, but also see Stevens et al., 2009). In contrast, most other regions of the semantic network showed at least some overlap between children and adults, especially in the left IFG, bilateral SMA, right insula, left MTG/STG, and left FG (see Figure 7C and Jackson, 2021).

Because meta-analyses depend critically on the quality of the underlying literature, they can be prone to a number of biases (e.g., positivity bias and selective reporting). Tools to detect the presence of such biases are less well developed in meta-analytic frameworks for neuroimaging as compared to clinical or behavioral outcomes (Acar et al., 2018). However, in the present study, the number of reported peak coordinates—a very rough analogue of an experiment-specific effect size for fMRI studies—was unrelated to the sample size of the experiments (see Figure 2). This is consistent with small sample bias and/or selective reporting in larger studies. To assess the robustness of our meta-analytic results against these kinds of biases, we first conducted a leave-one-out analysis which showed that all clusters were considerably invariant against the deletion of individual experiments from the sample. This means that we would have obtained identical results even if any of the original experiments reported only spurious activations. Second, we also conducted a fail-safe N analysis in which we estimated the number of unpublished null experiments (i.e., experiments without any consistent pattern of activation) that have to be added until the significance of any observed cluster is overturned. This number was larger than 30% of the meta-analytic sample size for almost all clusters. It thereby exceeded the current most conservative estimate for the actual number of unpublished fMRI experiments that are hidden in the “file drawer” (Samartsidis et al., 2020). Thus, although our meta-analysis could not directly assess or correct for publication bias in the underlying literature, its results seem to be stable even if one accepts that such biases are present.

## 5. Conclusion

Conducting fMRI experiments with children is challenging and costly, which is why sample sizes are often lower than in behavioral experiments or in neuroimaging experiments with adult participants. Meta-analyses are therefore necessary to filter out spurious results and to uncover similarities and differences between different task paradigms. Regarding children’s capacity to process semantic information, our coordinate-based meta-analysis showed reliable patterns of activation in the left IFG and MTG/STG, the bilateral SMA, the right insula and parts of the bilateral ventral temporal and occipital cortices. Within this network, tasks probing children’s semantic world knowledge and semantic relatedness between stimuli showed overlapping spots of activation that were distinct from those seen in tasks probing the differentiation of visual semantic object categories. A comparison to the adult semantic system revealed largely overlapping regions of activation but also more child-specific activation in bilateral inferior temporal and occipital regions as well as more adult-specific activation in the anterior portion of the left temporal lobe.

## Acknowledgements

We would like to thank the developers of the NiMARE software package (especially Taylor Salo and Tal Yarkoni) for their assistance as well as Rebecca L. Jackson, Timothy A. Keller, Saloni Krishnan, and Robin J. Schafer for providing us with additional data that were not publicly available. This research did not receive any specific grant from funding agencies in the public, commercial, or not-for-profit sectors.

## Appendix

Additional Information About the Experiments Included in the Meta-Analysis

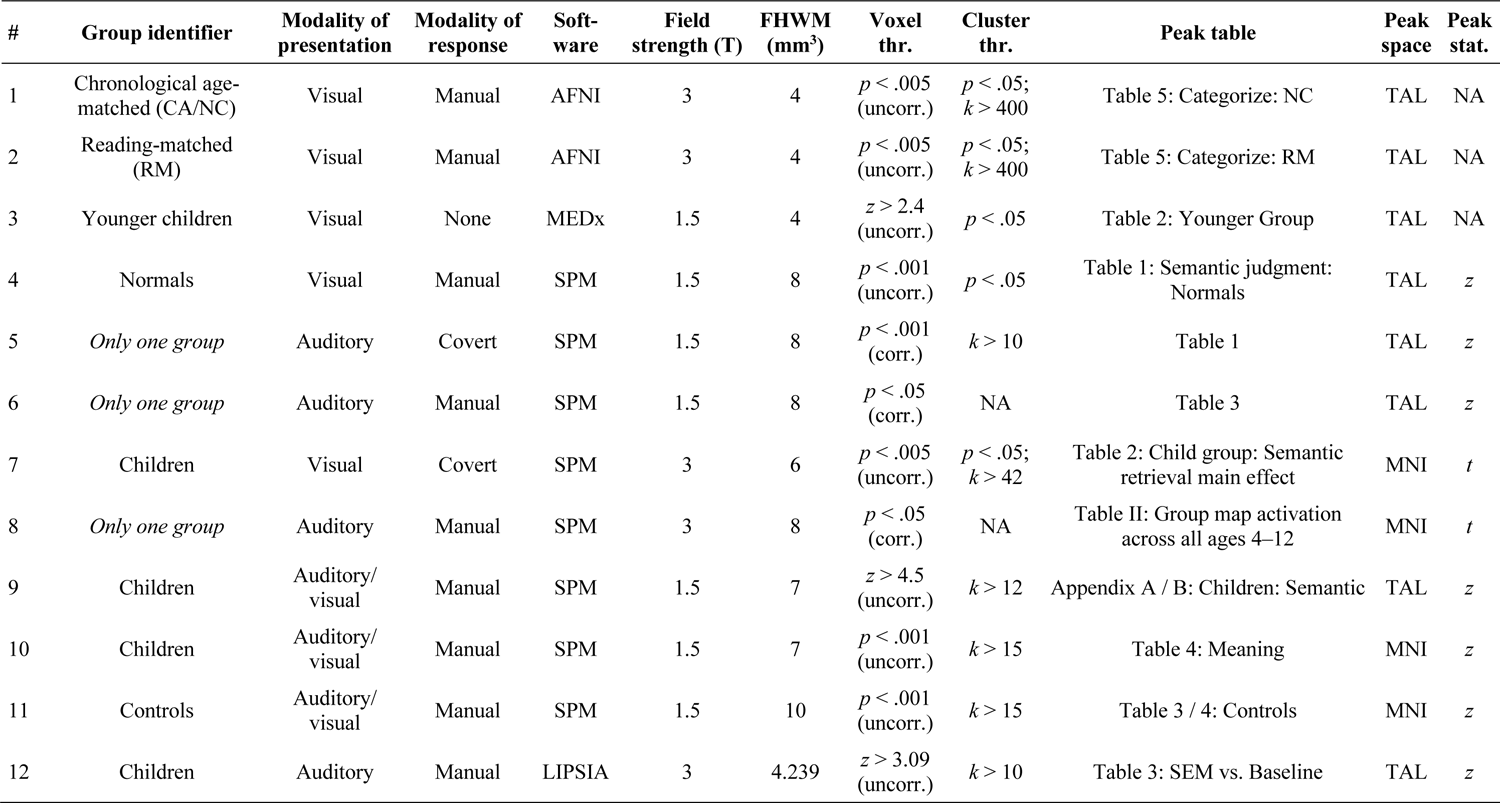
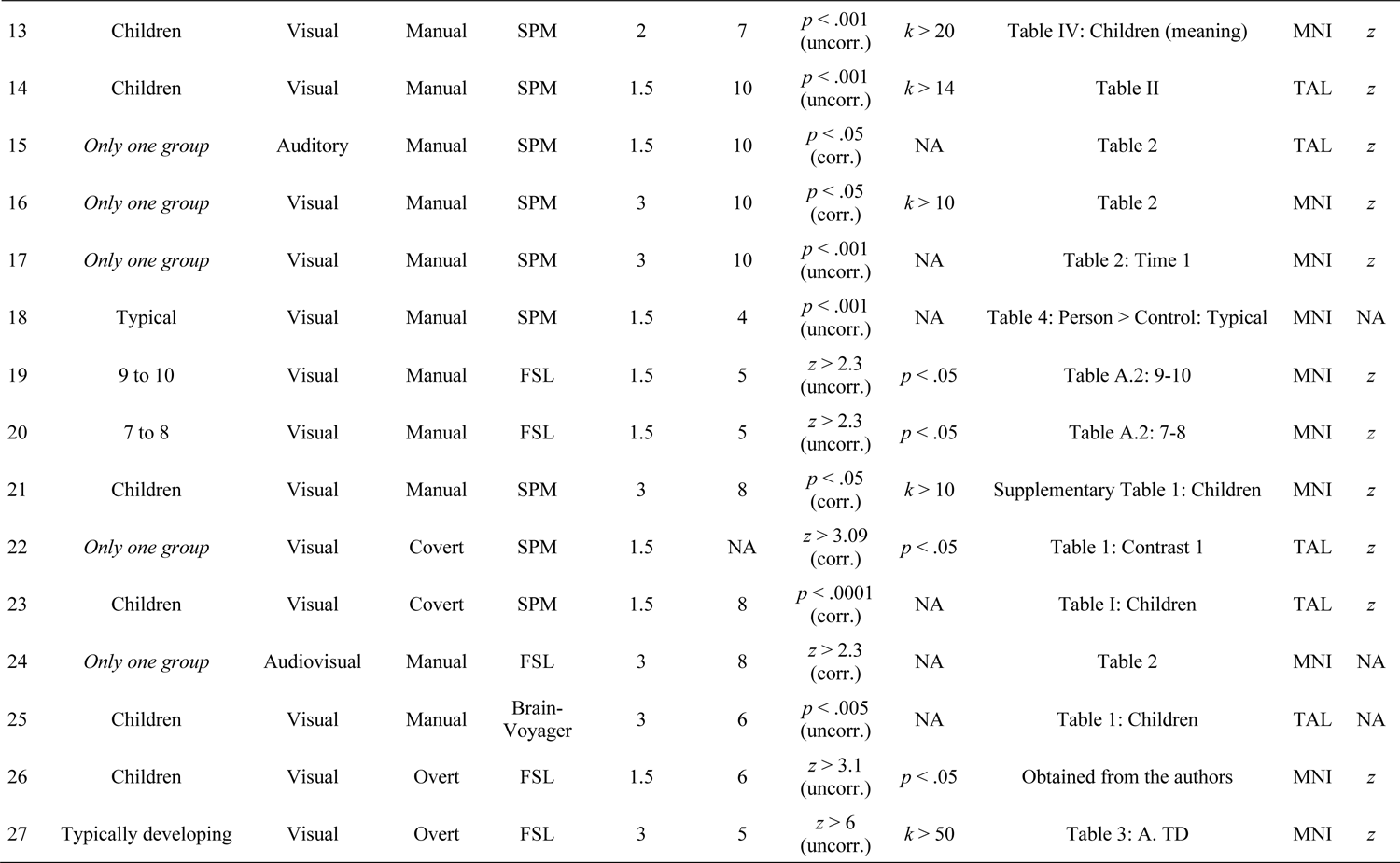
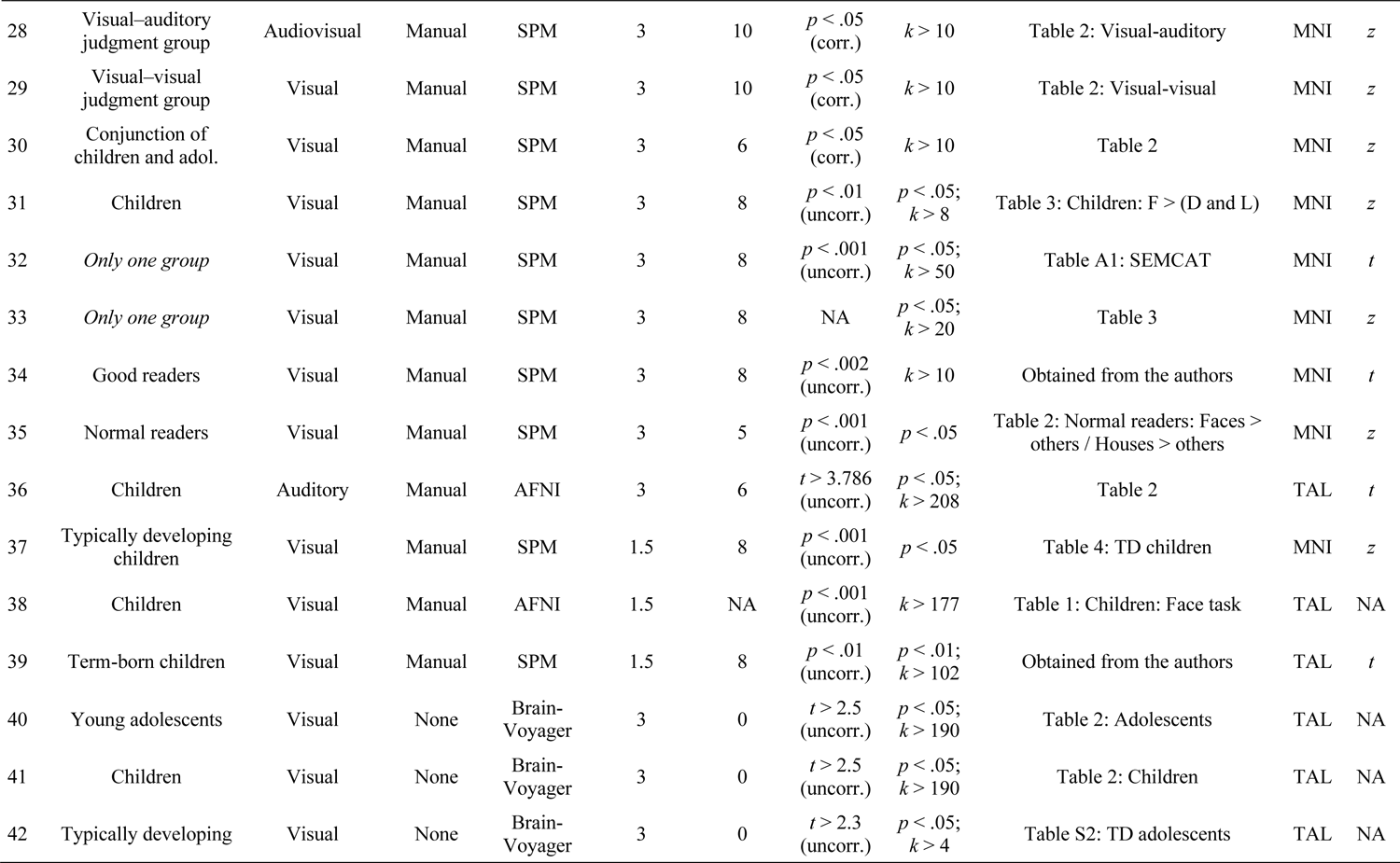
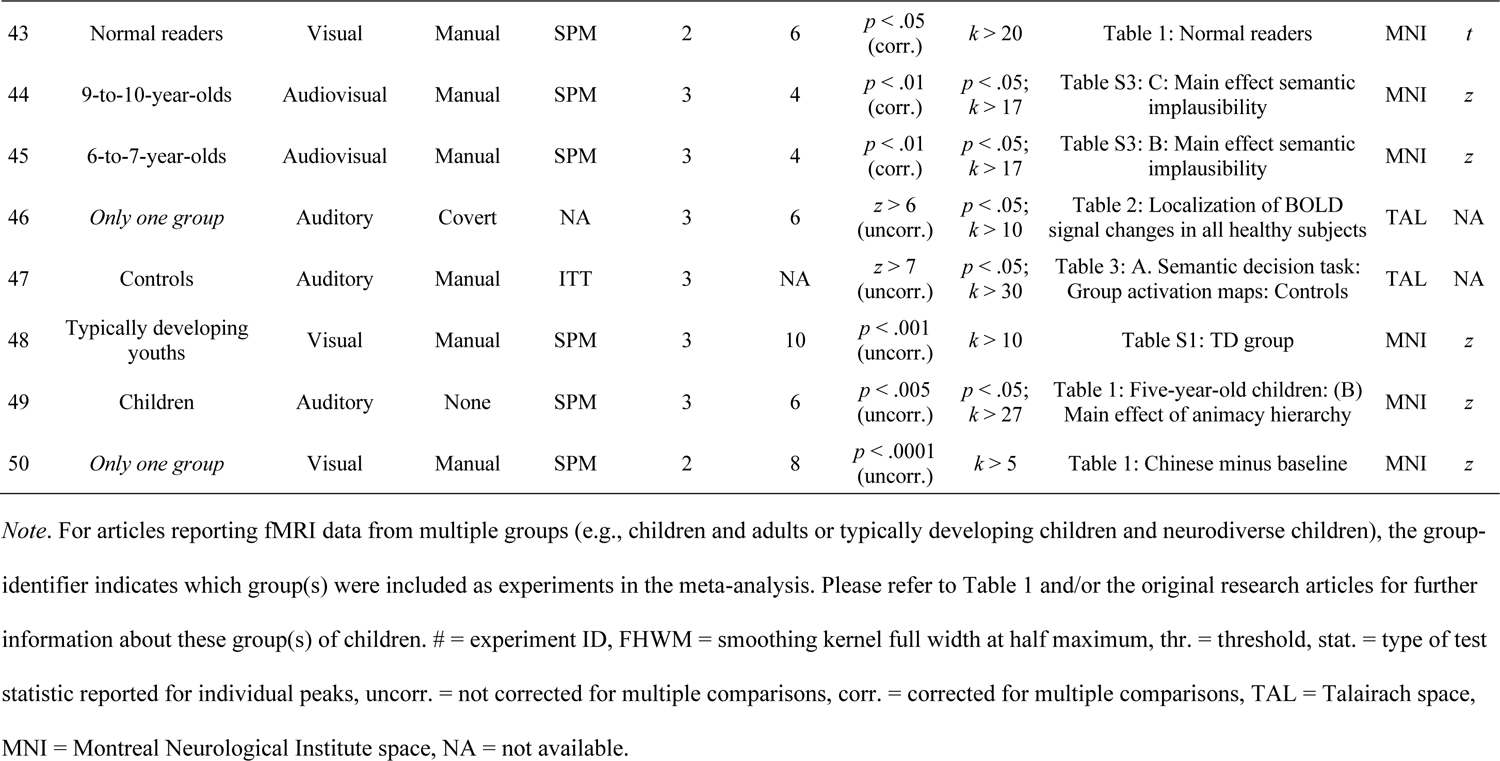

## Notes

https://osf.io/34ry2/

### Competing Interest Statement

The authors have declared no competing interest.

